# Hepatic Inactivation of Carnitine Palmitoyltransferase 1a Lowers Apolipoprotein B Containing Lipoproteins in Mice

**DOI:** 10.1101/2024.12.13.628437

**Authors:** Robert N. Helsley, Mikala M. Zelows, Victoria P. Noffsinger, Garrett B. Anspach, Nikitha Dharanipragada, Anna E. Mead, Isidoro Cobo, Abigail Carter, Qinglin Wu, Irina Shalaurova, Kai Saito, Josh M. Morganti, Scott M. Gordon, Gregory A. Graf

## Abstract

Genome- and epigenome-wide association studies have associated variants and methylation status of carnitine palmitoyltransferase 1a (CPT1a) to reductions in very low-density lipoprotein (VLDL) cholesterol and triglyceride levels. We report significant associations between the presence of *CPT1a* SNPs and reductions in plasma cholesterol, as well as positive associations between hepatic Cpt1a expression and plasma cholesterol levels across inbred mouse strains. Mechanistic studies show that both wild type and human apolipoprotein B100 (apoB)-transgenic mice with liver-specific deletion of *Cpt1a* (LKO) display lower circulating apoB levels consistent with reduced LDL-cholesterol (LDL-C) and LDL particle number. Despite a reduction in steady-state plasma lipids, VLDL-triglyceride (VLDL-TG) and cholesterol (VLDL-C) secretion rates are increased, suggesting accelerated clearance of apoB-containing lipoproteins (apoB-LPs) in LKO mice. Mechanistic approaches show greater peroxisome proliferator activated receptor α (PPARα) signaling which favors enhanced lipoprotein lipase-mediated metabolism of apoB-LPs, including increases in ApoCII and ApoAIV and reductions in ApoCIII & Angptl3. These studies provide mechanistic insight linking genetic variants and methylation status of *CPT1a* to reductions in circulating apoB-LPs in humans.

**HIGHLIGHTS:**

- Loss-of-function SNPs in *CPT1a* associate with reductions in plasma cholesterol in humans
- Hepatic Cpt1a expression positively associates with plasma cholesterol levels across inbred strains of mice
- Liver-specific Cpt1a deficiency lowers circulating apoB, plasma cholesterol, LDL-C, and LDL particle number
- Cpt1a ablation activates PPARα and favors clearance of apoB-containing lipoproteins

## INTRODUCTION

Dyslipidemia contributes to elevated risk of the manifestations of cardiometabolic disease including atherosclerotic cardiovascular disease, metabolic dysfunction-associated steatotic liver disease (MASLD), and type 2 diabetes. Familial hypercholesterolemia (FH) is estimated to affect 1 in 250-300 individuals worldwide (1). FH-causing variants in the low density lipoprotein receptor (LDLR), LDLR adaptor protein (LDLRAP), proprotein convertase subtilisin/kexin type 9 (PCSK9), apolipoprotein B (apoB), and APOE can be identified in 20-40% of individuals, indicating that the cause(s) of elevated LDL-cholesterol (LDL-C) is unknown in most patients (2). Thus, identification of new pharmacological targets that lower serum lipids (triglycerides, cholesterol) and improve metabolic disease outcomes are needed. One pathway that historically has been exploited for therapeutic benefit is that of mitochondrial fatty acid oxidation (FAO), with the notion that increasing the oxidation of long chain fatty acids may protect against facets of cardiometabolic disease. However, increasing mitochondrial FAO by targeting acetyl-CoA carboxylase (ACC) in patients with MASLD has been shown to increase apoB-LPs (3–11). Thus, the work herein aims to understand the counterintuitive association between hepatic long chain mitochondrial FAO and circulating apoB-LPs.

Carnitine palmitoyltransferase 1 (CPT1) is considered the rate-limiting enzyme for mitochondrial β-oxidation, where it is responsible for converting long chain acyl-CoAs to acyl-carnitines for entry into the mitochondria. The three isoenzymes of CPT1 (1a, 1b, 1c) differ in subcellular localization, catalytic activity, and tissue-specific expression patterns (12). For instance, CPT1a is highly expressed in the liver where it is sensitive to malonyl-CoA inhibition (13) and is critically important for fatty acid oxidation in response to fasting (14, 15). While the function of the CPT1b isoform is similar to that of CPT1a, it is most abundantly expressed in highly oxidative tissues such as heart, brown adipose tissue, and skeletal muscle (12, 16). The remaining isoform, CPT1c, is widely considered a brain-specific isoform that is expressed in the endoplasmic reticulum of neurons, binds to malonyl-CoA but does not support mitochondrial fatty acid oxidation (17–20).

Epigenetic (EWAS) and genome wide association studies (GWAS) have linked methylation sites within the proximal promoter and rare and common variants in CPT1a with a variety of traits associated with cardiometabolic disease including central obesity, adiposity and plasma triglycerides (21–23).

These associations are highly reproducible across multiple populations, suggesting a mechanistic link between CPT1a and cardiometabolic traits (24, 25). While variants that compromise CPT1a function would be expected to reduce FAO leading to increased storage in adipose tissue and accumulation in plasma, associations with steady-state levels of plasma cholesterol are less intuitive. Methylation of CpG islands near the *CPT1a* initiation codon reduce CPT1a mRNA and are negatively associated with plasma cholesterol, particularly among apoB-LPs (23). Further independent validation showed that *CPT1a* intron 1 was the lone site of hypomethylation in FH patients who have no underlying pathological variants in *LDLR*, *APOB*, or *PCSK9* (26). Taken together, these human association studies reveal a positive correlation between CPT1a expression and apoB-LPs. The goal of this work was to mechanistically explain these human associations by identifying the contribution of hepatic Cpt1a to lipid and lipoprotein phenotypes in wild type (WT) and human apoB100-transgenic (B100) mice.

## EXPERIMENTAL PROCEDURES

### Experimental Design and Study Subjects

Animal protocols were in accordance with NIH guidelines and approved by the Institutional Animal Care and Use Committee at the University of Kentucky. Mice were housed in individually ventilated cages at 20-22°C on a 14 h light/10 h dark cycle with ad libitum access to food and water. CPT1a-floxed (*Cpt1a*^F/F^) mice were obtained from the laboratory of Dr. Peter Carmeliet (27) and backcrossed >10 generations on the C57BL/6 background. To generate a mouse model exhibiting a more “humanized” lipoprotein profile, we crossed the human apoB100 (28, 29) and *Cpt1a*^F/F^ strains to generate B100/*Cpt1a*^F/F^ mice. Six to 8-week-old *Cpt1a*^F/F^ and B100/*Cpt1a*^F/F^ mice were intravenously administered 5×10^11^ adenoassociated virus (AAV) particles of either empty or AAVs encoding Cre-recombinase under control of a liver-specific promoter (TBG-Cre). The mice were then fed a semipurified high-fat diet (60% kcal from fat; Research Diets, D12492), low-fat diet (10% kcal from fat; Research Diets, D14042701), or western diet (WD; 42% kcal from fat; Research Diets, D12079B) for 12-16 weeks prior to necropsy. Mice were necropsied in the fasted (16-hour; 4 PM – 8 AM), fed (8 AM), or refed (16-hour fast followed by 6-hour refeed) state. Blood was collected by cardiac puncture, placed in EDTA-treated vials (VWR), and spun down at 10,000 x *g* for 10 minutes to collect plasma. Tissues were either collected and snap frozen in liquid nitrogen for biochemical assays or fixed in 4% paraformaldehyde for histological analysis.

Magnetic resonance relaxometry (Echo-MRI-100TM, Echo Medical System, Houston, TX) was used to assess total body composition. Serum glucose levels were measured using the INFINITY glucometer and glucose strips (ADW). Serum insulin (Crystal Chem) and β-hydroxybutyrate (Cayman Chem) levels were quantified using commercial assays according to manufacturers’ instructions.

### Histological Analysis

Hematoxylin and eosin (H&E) and picrosirius red staining of formalin-fixed paraffin-embedded liver sections were performed per manufacturers instructions and as previously described (15, 30–32).

### Assessment of bile acid and cholesterol secretion rates

Basal bile was collected as previously described with some modifications (33). Mice were anesthetized in a warmed chamber under 2% isoflurane, transferred to a Homeothermic Monitoring System (Harvard Apparatus #55-7030), and body temperature maintained at 37°C. A longitudinal, midline incision was made from the lower abdomen to the sternum. Two hemostatic forceps were used to separate the skin and expose the peritoneum. The peritoneum was incised and a wetted sterile cotton-tipped applicator used to expose the common bile duct and gall bladder. The bile duct was ligated with a 3 cm length of 3/0 silk suture. The gall bladder was cannulated with a 5-cm length of PE-10 tubing and secured with 3/0 silk suture. Bile was diverted into pre-weighed 0.5 ml microcentrifuge tubes and basal bile flow determined gravimetrically. Total biliary cholesterol and bile acid were determined using enzymatic colorimetric assays (Wako). Biliary lipid secretion rates are expressed as nmol/min/100g bw.

### Size exclusion chromatography of plasma lipoproteins

The lipoprotein cholesterol and triglyceride distributions were measured by size-exclusion chromatography, as previously described (34). Fifty to 200 μL of plasma were collected, diluted to a total volume of 500 μL in PBS, and loaded into an Akta Pure 25L liquid chromatography system.

Samples were passed through a Superose 6 Increase column (Cytiva) at a flow rate of 0.75 mL/min in PBS. Fractions were collected (0.5 mL/fraction) and assayed for cholesterol and triglycerides by enzymatic assays (Wako).

### Nuclear magnetic resonance (NMR) spectroscopy analysis of plasma lipoproteins

NMR was used to determine the size and composition of steady state plasma lipoproteins from B100^Con^ and B100^LKO^ mice. NMR spectra were acquired on a Vantera*^®^*Clinical Analyzer, a 400 MHz NMR instrument, from EDTA plasma samples as described for the NMR LipoProfile® test (Labcorp, Morrisville, NC) (35, 36). The NMR MetaboProfile analysis, using the LP4 lipoprotein profile deconvolution algorithm, reports lipoprotein particle concentrations and sizes. The diameters of the various lipoprotein classes and subclasses are: total triglyceride-rich lipoprotein particles (TRL-P) (24-240 nm), very large TRL-P (90-240 nm), large TRL-P (50-89 nm), medium TRL-P (37-49 nm), small TRL-P (30-36 nm), very small TRL-P (24-29 nm), total low density lipoprotein particles (LDL-P) (19-23 nm), large LDL-P (21.5-23 nm), medium LDL-P (20.5-21.4 nm), small LDL-P (19-20.4 nm), total high density lipoprotein particles (HDL-P) (7.4-13.0 nm), large HDL-P (10.3-13.0 nm), medium HDL-P (8.7-9.5 nm), and small HDL-P (7.4-7.8 nm). Mean TRL, LDL and HDL particle sizes are weighted averages derived from the sum of the diameters of each of the subclasses multiplied by the relative mass percentage. Linear regression against serum lipids measured chemically in an apparently healthy study population (n=698) provided the conversion factors to generate NMR-derived concentrations of total cholesterol (TC), triglycerides (TG), TRL-TG, TRL-C, LDL-C and HDL-C. NMR-derived concentrations of these parameters are highly correlated (r ≥0.95) with those measured by standard chemistry methods.

### Bile acid quantification by UPLC-MS/MS

Gall bladder bile was used for ultra-performance liquid chromatography-tandem mass spectrometry (UPLC-MS/MS) based quantification of bile acids in collaboration with the Biomarkers Core Laboratory at Columbia University, as previously described (37, 38).

### VLDL-TG & VLDL-C Secretion Rates

VLDL-TG and VLDL-C secretion rates were performed using previously established methods (39). Male and female B100^Con^ and B100^LKO^ mice were fed a western diet for 10-weeks. After 10-weeks of feeding, the mice underwent an overnight fast (16 h) prior to receiving 500 mg/kg of the nonionic detergent poloxamer 407 (P407), an inhibitor of LPL (39). Plasma was collected via tail vein at 2, 4, and 6 hours post P407 injection, and triglycerides and cholesterol were measured enzymatically (Wako).

### VLDL-TG Clearance

Frozen human plasma was obtained from the Kentucky Blood Center and lipoproteins isolated by sequential ultracentrifugation (40). Chylomicrons were floated by centrifugation (SW 32Ti, Beckman, 20K rpm, 30 min, 10° C) and removed by aspiration. Density was adjusted 1.019 g/ml with KBr, plasma centrifuged (VTi50, 55K rpm; 10°C, 12 hr), and VLDL isolated via tube cutter at the top of the clear zone. Density of the remaining plasma was adjusted to 1.21 g/ml with KBr, centrifuged (VTi50, 50K rpm; 10°C, 20 hr), and remaining lipoproteins removed via tube cutter to generate lipoprotein-deficient plasma (LPDP). VLDL and LPDP were extensively dialyzed against 150 mM NaCl, 0.01% EDTA (pH 7.4, 4oC, 14K MWCO, 4 × 4L), sterile-filtered through 0.45 μm filters (Millipore, Bedford, MA), and stored in aliquots under nitrogen at 4°C. To generate LPDS, LPDP was clotted with 1U/ml thrombin and centrifuged (Eppendorf, S-4-104, 3K rpm, 4oC, 15 min). Radiolabeled [^3^H]-triolein (NET431991MC Revvity, 600 mCi) was dried under streaming nitrogen in a borosilicate tube. VLDL (3 mg) was added to LPDS to a final volume of 3 ml as a source of cholesterol ester transfer protein, incubated overnight with agitation at 37°C, dialyzed as above, specific activity determined, and stored under nitrogen at 4°C.

Control (*n* = 6) and LKO (*n* = 5) mice were fasted (12 h), anesthetized (isoflurane, 2%), and radiolabeled VLDL (120 mCi) injected into the retro orbital sinus. Plasma (∼20 μl) was collected from the retro orbital sinus at 1, 5 and 10 min. Mice were exsanguinated via cardiopuncture at 15 min.

Radioactivity in plasma (10 μl) was determined by scintillation counting and clearance of VLDL expressed as percent of counts of 1 min following delivery of radiolabeled VLDL.

### Serum, hepatic, and fecal lipid analysis

Plasma and biliary cholesterol levels were measured using an enzymatic colorimetric assay (Wako). Triglycerides in plasma were quantified using similar methods (Wako). Absorbance was measured at 600 nm using the Biotek Synergy H1 Hybrid plate reader. Cholesterol was extracted from feces and analyzed by mass spectrometry as previously described (41–43).

Extraction of liver lipids and subsequent quantification of hepatic triglycerides, total and free cholesterol were conducted using enzymatic assays as previously described (15, 30–32). Initial liver mass was recorded then samples were delipidated in 4 mL of 2:1 chloroform to methanol (v/v) overnight. The organic solvent was dried under a constant stream of N_2_ prior to adding 6 mL of 2:1 chloroform to methanol. Dilute H_2_SO_4_ (0.05%) was added (1.2 mL), samples vortexed then centrifuged at 2000 rpm for 15 minutes to separate the phases. The bottom organic phase was recorded, and 0.5 mL of the organic phase was added to 1 mL of 1% TritonX-100 in chloroform. The samples were dried under N_2_ and 0.5 mL H_2_O was added prior to quantification of triglycerides (Wako), total and free cholesterol (Wako), per manufacturer’s instructions. All standards and blanks were prepared in a similar fashion, and data were expressed as μg lipid/mg of wet tissue.

### H3K27Ac ChIP-sequencing

#### Nuclei isolation from mouse liver tissue

Nuclei isolation was performed as described elsewhere (44, 45). Approximately 100 mg of frozen liver tissue was homogenized in 1 ml of 1% formaldehyde in Dulbecco’s phosphate-buffered saline (PBS; Corning) and incubated at room temperature for 10 minutes. The fixed tissue was quenched with 0.125 M glycine for 5 minutes at room temperature and pelleted at 1700 rpm for 10 minutes at room temperature. All subsequent steps were performed on ice or at 4°C. The tissue homogenate was washed three times with NF1 buffer containing 10 mM Tris-HCl pH 8.0, 1 mM EDTA, 5 mM MgCl2, 0.1 M sucrose, 0.5% Triton X-100 and incubated in 5 ml of NF1 buffer on ice for 30 minutes. Nuclei were extracted using a 7 ml Wheaton™ Dounce Tissue Grinder (DWK Life Sciences, 357542) and passed through a 70-μm strainer. The homogenate was underlaid with a 1.2 M sucrose cushion and centrifuged at 3,000 x g for 30 minutes. The pelleted nuclei were washed with NF1 buffer at 1,600 x g for 5 minutes, followed by two washes with a buffer containing (PBS, 1% bovine serum albumin, 1 mM EDTA) at 1,600 x g for 5 minutes. Nuclei for ChIP-Seq were stored at −80°C.

#### ChIP-seq library preparation and analysis

Chromatin immunoprecipitation (ChIP) was performed following previously described protocols (46, 47). Briefly, 1-2 million fixed nuclei from total liver tissue were thawed, pelleted by centrifugation, and lysed in 130 μl of LB3 lysis buffer containing 10 mM Tris-HCl (pH 7.5), 100 mM NaCl, 1 mM EDTA, 0.5 mM EGTA, 0.1% sodium deoxycholate, 0.5% N-lauroylsarcosine, 1x protease inhibitor cocktail, and 1 mM PMSF. Chromatin was fragmented by sonication using a PIXUL® Multi-Sample Sonicator in Corning™ 96-Well Clear Ultra-Low Attachment Microplates with the following settings: Pulse [N]: 50; PRF [kHz]: 1.00; Process Time: 60 minutes; Burst Rate [Hz]: 20.00. Protease inhibitors were freshly added to all buffers before use. The sonicated lysates were incubated overnight at 4°C with a mixture of Protein G Dynabeads (Invitrogen, #10003D) and H3K27ac antibodies (ActiveMotifs, #AB_2793305) on a rotator. The following day, immunocomplexes were captured using a magnet, and the beads were washed with 100 μl of pre-chilled buffers. Beads were sequentially washed four times with Wash Buffer I (20 mM Tris-HCl, pH 7.5, 150 mM NaCl, 1% Triton X-100, 0.1% SDS, 2 mM EDTA) and Wash Buffer III (10 mM Tris-HCl, pH 7.4, 250 mM LiCl, 1% Triton X-100, 0.7% sodium deoxycholate, 1 mM EDTA), followed by two washes with ice-cold TET buffer (10 mM Tris-HCl, pH 7.5, 1 mM EDTA, 0.2% Tween-20). Samples were transferred to a fresh PCR tube after the second wash to avoid the carryover of salts. ChIP libraries were prepared on beads using the NEBNext Ultra II Library Kit (NEB, #E7645L) with a 50% reduction in reaction volume, as described previously (48, 49). Reverse crosslinking was performed by adding 29 μl of a solution containing 4 μl 10% SDS, 3 μl 0.5 M EDTA, 1.6 μl 0.2 M EGTA, 1 μl 10 mg/ml Proteinase K (Biolabs, #P8107S), 1 μl 10 mg/ml RNase A plus 4.5 μl 5 M NaCl. Samples were incubated at 55°C for 1 hour, followed by 65°C overnight. After removing the Dynabeads, libraries were purified using 2 μl of SpeadBeads in 124 μl of 20% PEG 8000/1.5 M NaCl and washed twice with 100 μl of 80% ethanol.

The libraries were air-dried and eluted in 13 μl of buffer (10 mM Tris-HCl, pH 8.0, 0.05% Tween-20). PCR amplification was performed for 14 cycles using the NEBNext Ultra II PCR Master Mix with Solexa 1GA and 1GB primers. Finally, the libraries were size-selected (200-500 bp) on 10% TBE acrylamide gels (ThermoFisher, #EC62752BOX) and sequenced on a NovaSeq X-Plus.

H3K27ac ChIP-Seq peaks were aligned to open chromatin regions using the annotatePeak.pl tool from HOMER, with an ATAC-Seq peak file representing open chromatin regions of the mouse genome (50). Differential peaks were identified using HOMER’s getDiffExpression.pl tool with default parameters. DNA motifs within differential peaks or regions were identified using the findMotifGenome.pl tool with default settings, using random genomic sequences as background (51). De novo motifs are shown in **Figure 3**, unless specified otherwise.

### Real-Time PCR Analysis of Gene Expression

RNA extraction, cDNA synthesis, and quantitative real-time PCR was performed as previously described (15, 30–32). Frozen liver (∼20 mg) was homogenized in 1 mL of QIAzol (Qiagen 79306) and phase separated after the addition of 200 µL of chloroform. Approximately 400 μL of the aqueous phase was mixed with 400 µL of 75% ethanol prior to running the sample over a RNeasy spin column (Qiagen, RNeasy Mini Kit). 500 ng of RNA was used as a template to synthesize cDNA (High Capacity cDNA Reverse Transcription Kit, Applied Biosystems). QPCR reactions were carried out using 2X SYBR green (Cowin Biosciences) and mRNA expression levels were calculated using the ΔΔ-Ct method on an Applied Biosystems (ABI) QuantStudio 7 Flex Real-Time PCR System. Primers used for qPCR are listed in **Supplemental Table 1**.

### Bulk RNA-Sequencing

RNA was isolated from liver tissue using the RNeasy Mini kit (Qiagen). The quantity and quality of the samples were determined using the Cytation 5 (BioTek) plate reader and Agilent 4150 Tape Station System, respectively, prior to submission to Novogene. The mRNA-seq libraries were prepared and sequenced on an Illumina HiSeq2500 platform, at a depth of 20 M read pairs by Novogene. After sequencing, the paired-end clean reads were aligned to the reference genome using Hisat2 v2.0.5.

FeatureCounts v1.5.0-p3 was used to count the read numbers mapped to each gene, and the fragments per kilobase of transcript (FPKM) of each gene was calculated based on the length of the gene and read counts mapped to the gene.

Differential expression analysis was performed using the DESeq2 R package (1.20.0). The resulting p-values were adjusted using the Benjamini and Hochberg’s approach for controlling the false discovery rate. Genes with an adjusted p-value ≤0.05 found by DESeq2 were assigned as differentially expressed. Gene ontology (GO) enrichment analysis of differentially expressed genes was implemented by the clusterProfiler R package. GO terms with corrected p-values <0.05 were considered significantly enriched by differential expressed genes. The clusterProfiler R package was also used to test the statistical enrichment of differentially expressed genes in KEGG pathways (www.genome.jp/keg/).

### Liver single cell RNA-sequencing

#### Single cell suspensions

Liver single cell suspensions were obtained using reagents from the mouse liver dissociation and dead cell removal kits (Miltenyi Biotec). In short, the livers were excised from the animal, and caudal lobes and gallbladder were removed prior to cutting the liver into small tissue pieces (∼4 pieces of liver per lobe [right, left, medial]). All tissue pieces were placed into tube C containing 4.7 mL of DMEM/F12 (Corning) containing 200μL of enzyme D, 100μL enzyme R, and 20μL of enzyme A. Tube C was then placed in the gentleMACS^TM^ Octo Dissociator with Heaters and digested at 37°C for 30 minutes. The homogenate was then ran over a 70μm strainer and rinsed with 5mL of cold DMEM/F12. The cells were centrifuged at 300 x *g* for 10 minutes, and the cell pellet was resuspended in 4 mL of debris removal solution then overlaid with 4 mL of 1X Dulbecco’s Phosphate-Buffered Saline (DPBS). After another 3000 x *g* spin, the supernatant was removed and the cell pellet was resuspended in 5mL of 1X red blood cell (RBC; eBioscience^TM^) lysis buffer. After 5 minutes, the RBC lysis buffer was neutralized with 9mL of DMEM/F12 then centrifuged at 300 x *g* for 5 minutes at 4°C. The cell pellet was then resuspended in 500μL of 1X DPBS + 1% BSA and ran over a 70μm Flowmi cell strainer (twice) before adding ∼4.5mL of fresh DPBS + 1% BSA prior to cell counting and library preparation.

#### Library preparation and sequencing

Cell suspensions were diluted to a target concentration of approximately 1,000 cells/μL in PBS + 1% BSA. Cells were counted by hemocytometer and the fraction of live verse dead cells was determined using acridine orange and ethidium homodimer. Cells were loaded into wells of a 10x Genomics Next GEM Chip G to target the capture of 10,000 cells per sample. Reverse transcription, amplification, and library production were performed using the 10x Genomics 3’ Gene Expression Next GEM Kit v3.1 according to the manufacturer’s instructions (10x Genomics, User Guide CG000315). Libraries were sequenced at the University of Kentucky Genomics Core Laboratory on a NovaSeq 6000 S4 flow cell to a depth of about 20,000 read pairs per cell.

#### Data analysis

Sequence alignment, cell calling, and initial clustering was performed using default parameters with Cell Ranger v7.0.0 (10x Genomics) on the University of Kentucky HPC Morgan Compute Cluster. The pre-built mouse mm10-2020-A reference transcriptome (downloaded from 10x Genomics) was used. Standard pre-processing protocols were followed in Cell Ranger to identify cells above background and the resulting filtered gene matrices were utilized in Seurat (v4.0.3) for single cell analyses (52, 53). Genes that were detected in 3 or more cells were used for analyses. Single cell QC was performed to exclude cells with less than 200 genes or those that exceeded 2500 genes, and greater than 25% mitochondrial genes. The three biological replicates per genotype were merged into a single Seurat object. Data were normalized, scaled, and variable features found using *SCTransform* with default parameters for the merged object. Subsequently, PCA and UMAP were used for two-dimensional visualization of the dimensionally reduced dataset (with the top 30 PCs used, based upon the total variance explained by each). *FindAllMarkers* function was used to identify the top10 differentially expressed genes for each cell cluster, cells clusters expressing canonical markers were used to define main classes from liver tissues, as shown previously (54, 55). Pairwise contrasts were extracted using *FindMarkers* function in tandem with *MAST* for calculating differentially expressed genes (DEGs) as a function of genotype (i.e. LKO vs. WT). DEGs were plotted using EnhancedVolcano package (56).

### Immunoblotting

Whole liver (∼20 mg) homogenates were solubilized in 1 mL of 1X ice-cold RIPA buffer (Cell Signaling) supplemented with 1X Halt Protease and Phosphatase Inhibitor Cocktail (ThermoFisher). After two sequential centrifugation steps (13,000 rpm for 10 min), the supernatant was collected and protein was quantified using the Pierce^TM^ Bicinchoninic Acid Protein (BCA) assay (ThermoFisher). For SDS-PAGE, 10 µg of protein was loaded and separated on 4-15% criterion TGX gels. Proteins were blocked with 5% milk and probed with antibodies listed in **Supplemental Table 2**. Images were taken on a ChemiDoc MP Imaging System (Bio-Rad) and quantification of blots were performed using ImageJ software (NIH).

### Statistical Analysis

All graphs and statistical analyses were completed using GraphPad Prism 10.2.3. Data are expressed as ± SEM, unless otherwise noted in the figure legends. Two-way ANOVAs followed by a Tukey’s multiple comparison *post hoc* analysis was used when comparing two variables (e.g., genotype x diet). To assess differences between endpoints which include time as a variable (e.g., body weights, IPGTT, caloric intake, etc.), a repeated measures two-way ANOVA was performed. When comparing two groups, a Student’s two tailed *t*-test was utilized. P-values <0.05 were considered statistically significant. All figure legends contain the statistical analysis used for each panel of data.

## RESULTS

### Gene Expression and Genetic Variants of *CPT1a* Associate with Plasma Cholesterol Levels in Mice and Humans

Previous reports have identified strong associations of *CPT1a* variants with plasma lipid traits across independent human cohorts (26, 57–59). We next asked whether SNPs within *CPT1a* also associated with plasma lipid traits across human populations. Utilizing the Common Metabolic Diseases Knowledge Portal, we determined which individual SNPs were most highly associated with plasma cholesterol levels (total and LDL-C). Intriguingly, the top 10 most highly significant SNPs based off *P*-value all had negative associations with both total cholesterol (**Figure 1A**) and LDL-C (**Figure 1B**). The top SNP (based off *P*-value) that negatively associated with plasma cholesterol levels was rs2229738, a missense variant (A275T) shown to decrease CPT1a enzymatic activity by 40% (60).

**Figure 1.**
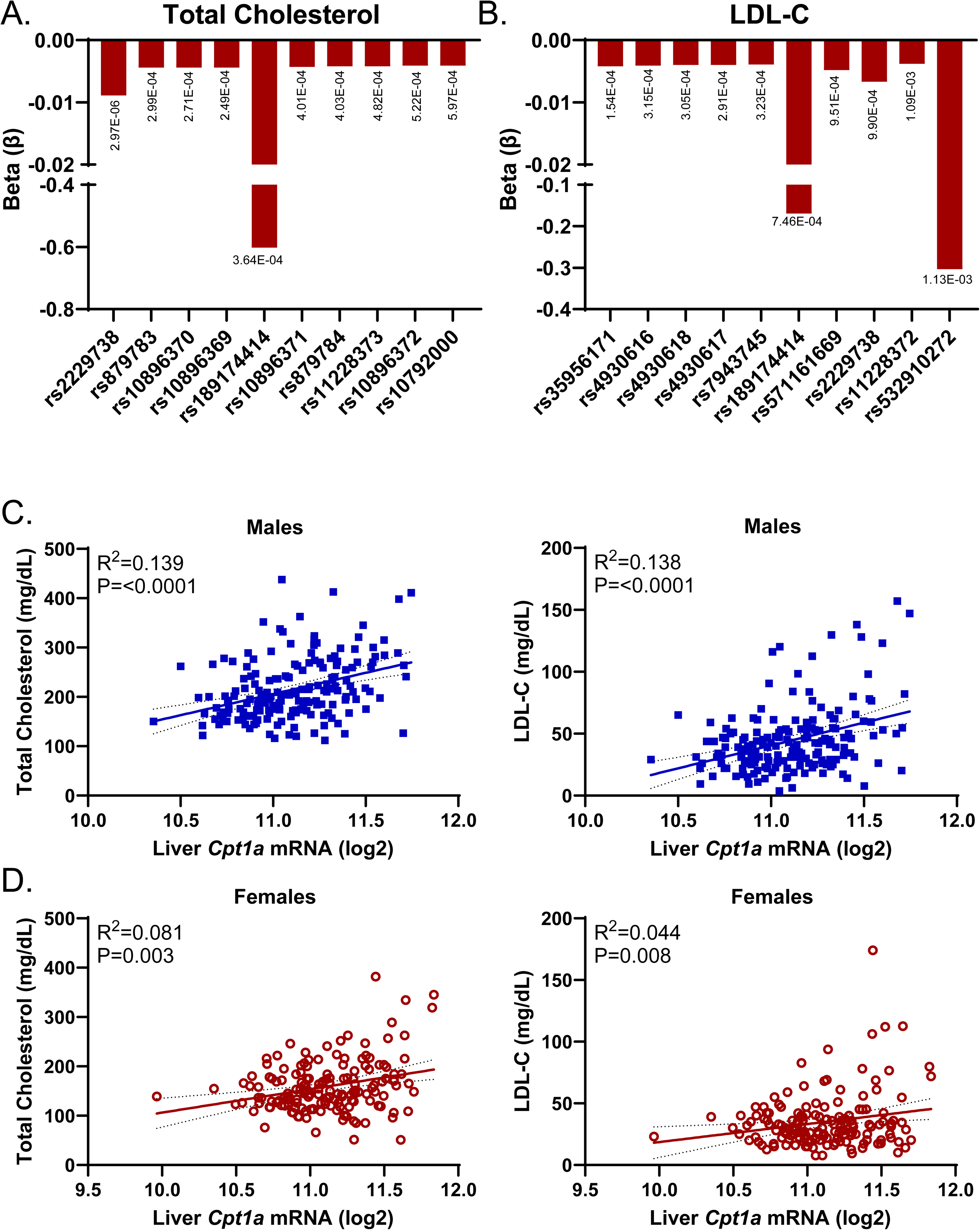
*CPT1a* SNPs and RNA Levels Correlate with Total Cholesterol and LDL-C. (**A, B**) The directionality and effect size (β) of the top 10 most highly significant SNP-cholesterol associations in CPT1a. Total cholesterol (**A**) and LDL-C (**B**). Data presented from **A** and **B** were obtained from the Common Metabolic Diseases Knowledge Portal (CMDKP). (**C-D**) Linear regression analysis of hepatic Cpt1a RNA levels with total cholesterol (left) and LDL-C (right) in males (**C**) and females (**D**) across inbred strains of mice from the hybrid mouse diversity panel (n=158-182). All *P*-values are listed in the figure.

We then utilized a systems genetics approach and leveraged data from the hybrid mouse diversity panel (HMDP) to examine links between Cpt1a expression and plasma total cholesterol and LDL-C (61). To induce obesity, ∼100 inbred strains of mice were *ad libitum* fed a high fat, high sucrose diet for 8-weeks prior to plasma lipid measurements (62). Consistent with prior human associations, hepatic Cpt1a gene expression positively associated with total cholesterol and LDL-C in male (*p*<0.0001; **Figure 1C**) and female (*p*<0.003 total; *p*<0.008 LDL-C; **Figure 1D**) mice. Taken together, these data implicate CPT1a as a potential modulator of plasma cholesterol phenotypes in mice and humans.

### Liver-specific Cpt1a Deletion Reduces Circulating Cholesterol Levels in WT Mice

Given the strong associations of *CPT1a* variants with plasma lipid traits across independent human cohorts (**Figure 1A, B**) (26, 57–59) and inbred mouse strains (**Figure 1C, D**), we first sought to determine the contribution of hepatic Cpt1a to lipid and lipoprotein metabolism in wild-type mice. Male *Cpt1a*^F/F^ mice were intravenously injected with AAVs containing empty (Control) or TBG-Cre to drive liver-specific (63) deletion (LKO) of Cpt1a (**Figure 2A**, **B**). Mice were then fed a high fat diet (HFD; 60% kcal from fat) for an additional 12-weeks prior to a fasting (16h) and refeeding (16h fast followed by 6h refeed) regimen at necropsy. An experimental overview is provided in **Supplemental Figure 1A**. Despite similar caloric intake, male LKO mice exhibited a slight, albeit significant (genotype x diet interaction: *p*<0.0001), decrease in body weight throughout HFD-feeding (**Supplemental Figure 1B**).

**Figure 2.**
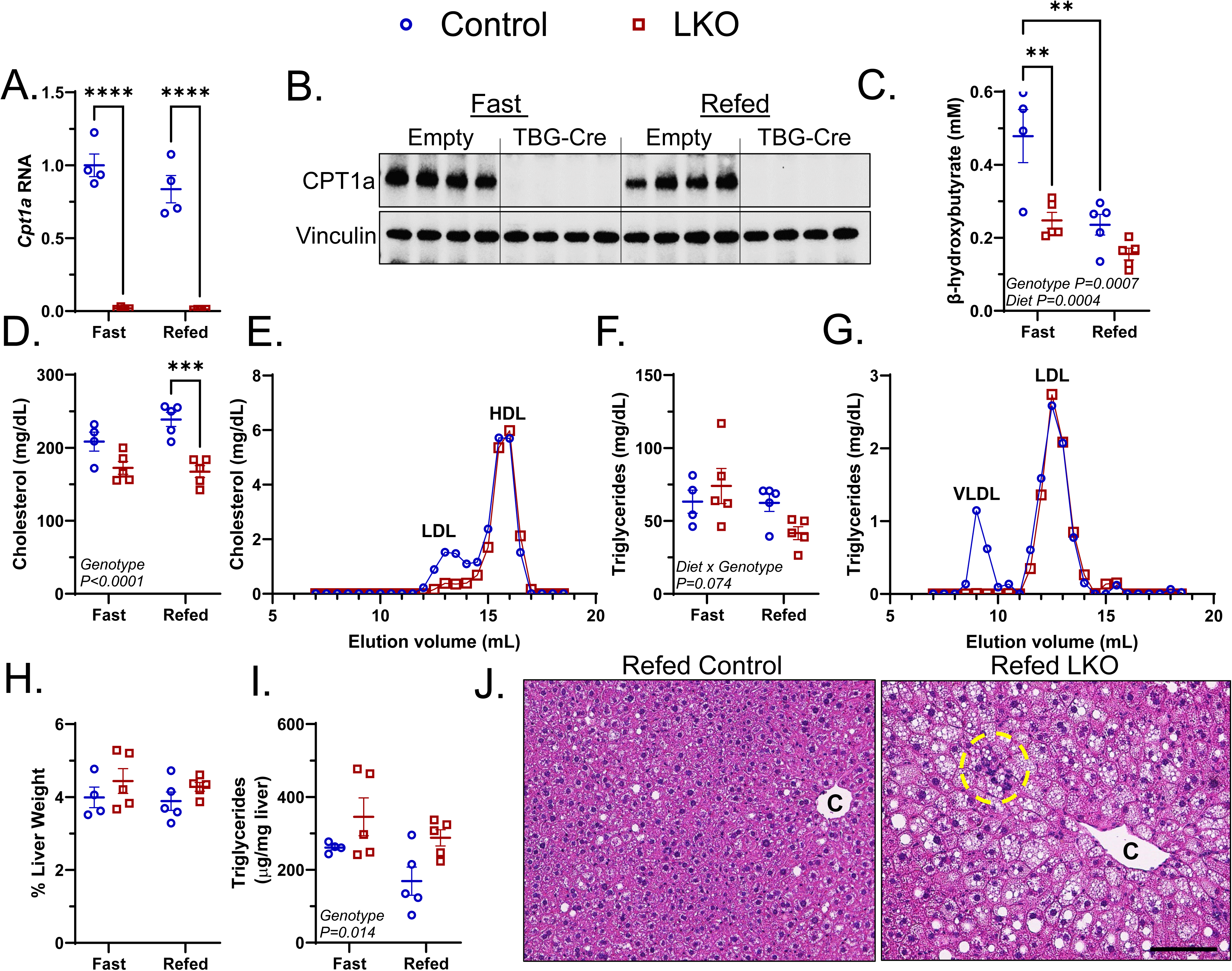
Liver-specific Cpt1a Deletion Reduces Plasma Cholesterol Levels in WT Mice. Male *Cpt1a^F/F^* mice were treated with AAVs containing empty or TBG-Cre prior to HFD-feeding for 12 weeks. Mice were necropsied in the fasted or refed state. (**A-B**) Liver Cpt1a RNA and protein levels were measured by qPCR (**A**; n=4) and western blot (**B**; n=4). Vinculin is a loading control. (**C**) β-hydroxybutyrate levels were measured from plasma of fasted and refed mice (n=4-5). (**D-G**) Steady-state cholesterol and triglycerides in whole plasma (**D**, **F**) and in FPLC-fractionated plasma (**E**, **G**) from WT and LKO mice. (**H-J**) Liver weight normalized to body weight (**H**), hepatic triglycerides (**I**), and liver H&E (**J**) staining from control and LKO mice (n=4-5). Significance was determined by two-way ANOVA with Tukey’s multiple comparison *post hoc* analysis. **P<0.01; ***P<0.001; ****P<0.0001.

To confirm successful deletion of Cpt1a, we measured Cpt1a RNA and protein levels in the liver by real-Time PCR (qPCR) and immunoblotting, respectively. This loss-of-function approach reduced Cpt1a RNA and protein levels by greater than 95% in LKO mice across both dietary regiments (**Fig. 2A, B**). We then measured plasma β-hydroxybutyrate levels as a surrogate readout of hepatic FAO. We observed significant reductions in circulating β-hydroxybutyrate levels in LKO mice regardless of nutritional status (significance by genotype; *p*=0.0007); however, the observed difference between control and LKO mice was diminished with refeeding (**Fig. 2C**).

Primary manifestations associated with CPT1a deficiency include hypoketotic hypoglycemia (64). Consistent with these clinical manifestations, mice with liver-specific Cpt1a deletion showed a 30% reduction in fasting blood glucose levels with no observable differences upon refeeding (**Supplemental Figure 2A**). Further, *Cpt1a* deletion resulted in a ∼75% reduction in fasting insulin levels and homeostatic model assessment for insulin resistance (HOMA-IR), while exhibiting improved insulin sensitivity as measured by intraperitoneal insulin tolerance tests (**Supplemental Figure 2B-E**). However, genotypic differences were not observed in intraperitoneal glucose tolerance tests.

We next assessed whether Cpt1a deletion in the liver would influence lipoprotein profiles and hepatic lipids in response to fasting and refeeding. We observed a ∼30% reduction in plasma cholesterol while triglycerides approached significance (*P*=0.074) with refeeding in LKO mice (**Fig. 2D, F**). Size exclusion chromatography on plasma from refed mice revealed significant reductions in LDL-C (**Fig. 2E**) and VLDL-TGs (**Fig. 2G**), with no observable differences in high density lipoprotein cholesterol (HDL-C). Further, no changes in liver mass were observed (**Fig. 2H**) but LKO led to modest increases in hepatic triglycerides, largely with refeeding (significance by genotype=0.014; **Fig. 2I**).

Hematoxylin and eosin staining revealed more abundant accumulation of lipid droplets and inflammatory cells in refed LKO mice as compared to controls (**Fig. 2J**). Taken together, male LKO mice exhibit reductions in plasma cholesterol and triglycerides with modest increases in hepatic triglyceride levels.

### Liver-specific Cpt1a Deletion Activates PPARα Signaling & Alters Apolipoprotein Levels

In an attempt to elucidate the mechanistic underpinnings linking hepatic Cpt1a to lower plasma lipids, we first measured nuclear protein levels of transcription factors known to be affected by fasting and refeeding. As anticipated, nuclear levels of the sterol response element binding protein 1 (SREBP1) and the carbohydrate response element binding protein (ChREBP) were significantly elevated in control mice with refeeding (**Fig. 3A**). Peroxisome proliferator activated receptor α (PPARα), which mediates the transcriptional response to fasting (65), was repressed in male control mice during refeeding (**Fig. 3A**). In response to Cpt1a deletion, male mice were unable to upregulate SREBP1 but nuclear ChREBP levels were elevated in LKO mice relative to controls during refeeding. Nuclear PPARα levels were greater in LKO mice regardless of nutritional status and repressed to similar degrees in both genotypes with refeeding (**Fig. 3A**).

**Figure 3.**
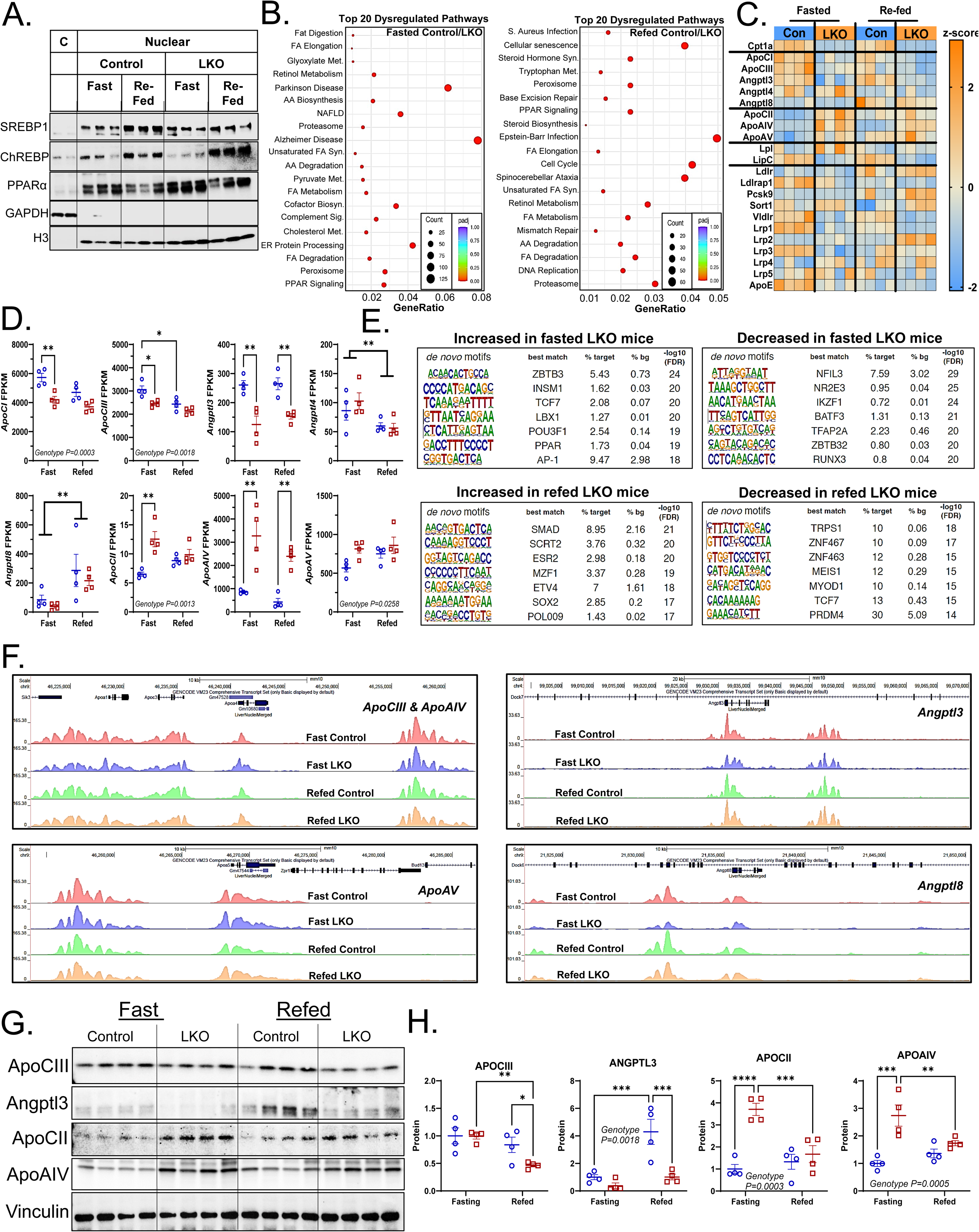
Liver-specific Cpt1a Deletion Activates PPARα and Alters H3K27 Acetylation. Male *Cpt1a^F/F^* mice were treated with AAVs containing empty or TBG-Cre prior to HFD-feeding for 12 weeks. Mice were necropsied in the fasted or refed state. (**A**) Cytoplasmic (“C”) and nuclear protein levels of SREBP1, ChREBP, and PPARα. GAPDH and H3 were used as loading controls for cytoplasmic and nuclear protein, respectively (n=3). (**B-D**) Bulk RNA-sequencing was performed on livers from control and LKO mice in the fasted and refed state. KEGG enrichment dot plots (**B**), heatmaps with z-scores spanning −2 to +2 (**C**), and individual expression of genes known to modulate LPL activity (**D**). Data are presented as fragments per kilobase of transcript per million mapped reads (FPKM). (**E**) *De novo* DNA motifs from H3K27-Ac ChIP-sequencing that are increased or decreased in LKO mice upon fasting and refeeding. (**F**) H3K27-Ac ChIP-sequencing peaks aligned to chromatin regions (*ApoCIII, ApoAIV, ApoAV, Angptl3, Angptl8*) using the USCS genome browser. (**G-H**) Immunoblotting (**G**) and subsequent densitometry (**H**) of ApoCIII, Angptl3, ApoCII, and ApoAIV, proteins identified as being differentially regulated in **D**. Vinculin serves as a loading control. Significance was determined by two-way ANOVA with Tukey’s multiple comparison *post hoc* analysis. *P<0.05; **P<0.01; ***P<0.001; ****P<0.0001.

To complement the transcription factor analysis, we completed bulk RNA-sequencing on liver tissue from control and LKO fasted and refed mice. In the fasted state, we identified 969 genes that met pre-determined statistical (*p*-adjusted value ≤0.05) and effect size cut-offs (Log2 normalized fold-change [FC] of ±0.58). Of the 969 genes identified, 444 were upregulated while 525 were downregulated in response to *Cpt1a* deletion. The most significantly downregulated genes in male LKO mice were *Srebf1* (Log2 FC=-2.26; Padj=1.34E-50) followed by *Cpt1a* (Log2 FC=-2.28; Padj=1.35E-40). KEGG enrichment analysis revealed that pathways involved in the biosynthesis of amino acids (*Pklr, Glul, Pah*) and protein processing in the endoplasmic reticulum (*Sec61b*, *Ddost*, *Hyou1*; **Fig. 3B**) were significantly downregulated in LKO compared to control mice. On the contrary, the most significantly upregulated genes in fasted male LKO were laminin subunit beta 3 (*Lamb3*; Log2 FC=2.10; Padj=6.07E-33) and fatty acid elongase 5 (*Elovl5*; Log2 FC=1.76; Padj=2.46E-32), consistent with greater enrichment of PPAR signaling genes (**Fig. 3B**) and elevations in nuclear PPAR protein levels (**Fig. 3A**). In the refed state, 546 genes were identified with 340 upregulated and 206 downregulated in response to *Cpt1a* deletion. The most significantly downregulated genes in refed male LKO mice were *Cpt1a* (Log2 FC=-3.06; Padj=1.35E-34) followed by *Rcan2* (Log2 FC=-2.58; Padj=1.01E-19), while pathways involved in complement and coagulation cascades (*Plg*, *Cpb2*, *C3*) and retinol metabolism (*Ugt2b1, Cyp2a5, Cyp2c50*; **Fig. 3B**) were significantly downregulated in LKO compared to control mice. The most significantly upregulated genes in refed male LKO were immunoglobulin kappa constant (*Igkc*; Log2 FC=4.03; Padj=2.63E-52) and enoyl-Coenzyme A hydratase (*Ehhadh*; Log2 FC=2.42; Padj=8.82E-21), consistent with greater immune cell infiltration (**Fig. 2J**) and activation of PPAR signaling, respectively (**Fig. 3B**). Consistent with an increase in nuclear PPARα protein and enrichment of PPARα-target genes by bulk RNA-sequencing, proteins that promote mitochondrial (LCAD, SCAD) and peroxisomal (PMP70, ACOX1) β-oxidation were elevated by 1.5 to 2 fold in response to *Cpt1a* deletion (**Supplemental Figure 3A, B**).

One of the clinical benefits of PPARα agonist treatment is the lowering of serum cholesterol and triglycerides levels (66). A mechanism by which fibrates lowers apoB-LPs is through repression of the apolipoprotein C-III gene, which leads to enhanced lipoprotein lipase (LPL)-mediated catabolism of these particles (66–68). Therefore, we examined normalized reads of individual genes known to be regulated by PPARα that influence lipoprotein metabolism. We observed significant reductions in genes known to inhibit LPL activity (*ApoCI*, *ApoCIII*, *Angptl3*) while genes known to enhance LPL activity (*ApoCII*, *ApoAIV*, *ApoAV*) were largely increased in LKO mice (**Fig. 3C, D**). While these gene expression changes are consistent with that of PPARα activation, we sought to analyze the enhancer landscape by measuring H3K27 acetylation (H3K27ac), an epigenetic mark of active enhancers (69).

Chromatin immunoprecipitation-sequencing (ChIP-seq) studies revealed the top H3K27ac-enriched DNA motifs were zinc finger and BTB domain containing 3 (Zbtb3) and mothers against decapentaplegic homolog (Smad) in fasted and refed LKO mice, respectively (**Fig. 3E**). Notably, the *de novo* motif for PPAR was enriched with H3K27ac in fasted LKO mice (**Fig. 3E**). Further, the top motifs associated with H3K27ac enrichment that were decreased in fasted and refed LKO mice were the nuclear factor, interleukin 3 regulated (NFIL3) and the transcriptional repressor GATA binding 1 (TRPS1; **Fig. 3E**). H3K27ac ChIP-Seq peaks were then aligned to open chromatin regions of apolipoprotein loci that were differentially altered by RNA-sequencing. The amplitude of the HK37ac ChIP-peak at the *ApoCIII* locus was slightly reduced in both fasted and refed LKO mice, while there appears to be a shift in the localization of the HK37ac ChIP-peak within the *ApoAIV* locus (**Fig. 3F**).

Further, the amplitude of the HK37ac ChIP-peaks within the *ApoAV* locus was greater in the LKO mice, while the peak at the *Angptl3* locus were considerably reduced in LKO mice (**Fig. 3F**).

We next determined if gene expression changes were reflected at the protein level by immunoblot analysis. We observed a 30% reduction in ApoCIII in the refed state and a 50-75% reduction in Angptl3 irrespective of nutritional status. Conversely, ApoAIV and ApoCII protein increased 3-4 fold in LKO mice in the fasted state (**Fig. 3G, H**). Taken together, liver-specific deletion of Cpt1a in male mice increases PPARα- and H3K27ac-driven transcriptional responses that favor accelerated LPL-mediated clearance of apoB-LPs.

### Liver-specific Cpt1a Deletion Lowers Plasma Cholesterol and Triglycerides in Human ApoB100-transgenic Mice

One major limitation to studying lipoprotein metabolism in rodents, particularly of apoB-LPs, is that mice carry most of their cholesterol in HDL as opposed to humans which carry most of their cholesterol in LDL. Therefore, we next sought to determine if inactivation of *Cpt1a* would reduce apoB-LPs in a mouse model with a “humanized” lipoprotein profile. To do this, we bred *Cpt1a*^F/F^ and human apoB100 transgenic (28, 29) mice to generate a B100/*Cpt1a*^F/F^ strain. We first validated the model and showed that human apoB100 mRNA and protein were only detected in the liver and plasma, respectively, from B100/*Cpt1a*^F/F^ mice (**Supplemental Figure 4A, B**). Similar to human lipoprotein profiles, LDL-C is the predominant carrier of plasma cholesterol in B100/*Cpt1a*^F/F^ mice with no observable difference in HDL-C (**Supplemental Figure 4C**).

Next, B100/*Cpt1a*^F/F^ mice received an intravenous injection of either empty AAV or TBG-Cre AAV (similar approach to **Figure 1**) to yield B100 control (B100^Con^) or B100 CPT1a LKO (B100^LKO^) mice. The B100^Con^ and B100^LKO^ mice were then fed a semipurified low fat (LFD) or western diet (WD) for 16 weeks prior to necropsy. Feeding a WD increased body weight and percent adiposity in both male and female mice, while loss of CPT1a in the liver increased body weight and adiposity levels only in female mice (**Supplemental Figure 5A, B**). Similar to Figure 1, AAV-TBG treatment reduced CPT1a RNA and protein levels in the liver by greater than 90% in LKO mice fed both a LFD and WD (**Fig. 4A, B**). Male B100^Con^ mice fed a LFD and WD had steady-state plasma triglyceride levels of 226±43 and 249±98 mg/dL, while male B100^LKO^ mice had reduced circulating triglyceride levels of 96±31 and 116±25 mg/dL when fed LFD and WD, respectively (**Fig. 4C**). Similarly, female B100^Con^ mice exhibited plasma triglyceride levels of 132±35 and 82±25 mg/dL, while B100^LKO^ mice had levels of 81±26 and 43±18 mg/dL when maintained on LFD and WD diets, respectively (**Fig. 4C**). Overall, male and female B100^LKO^ mice had ∼40-60% reductions in steady-state triglyceride levels irrespective of diet.

**Figure 4.**
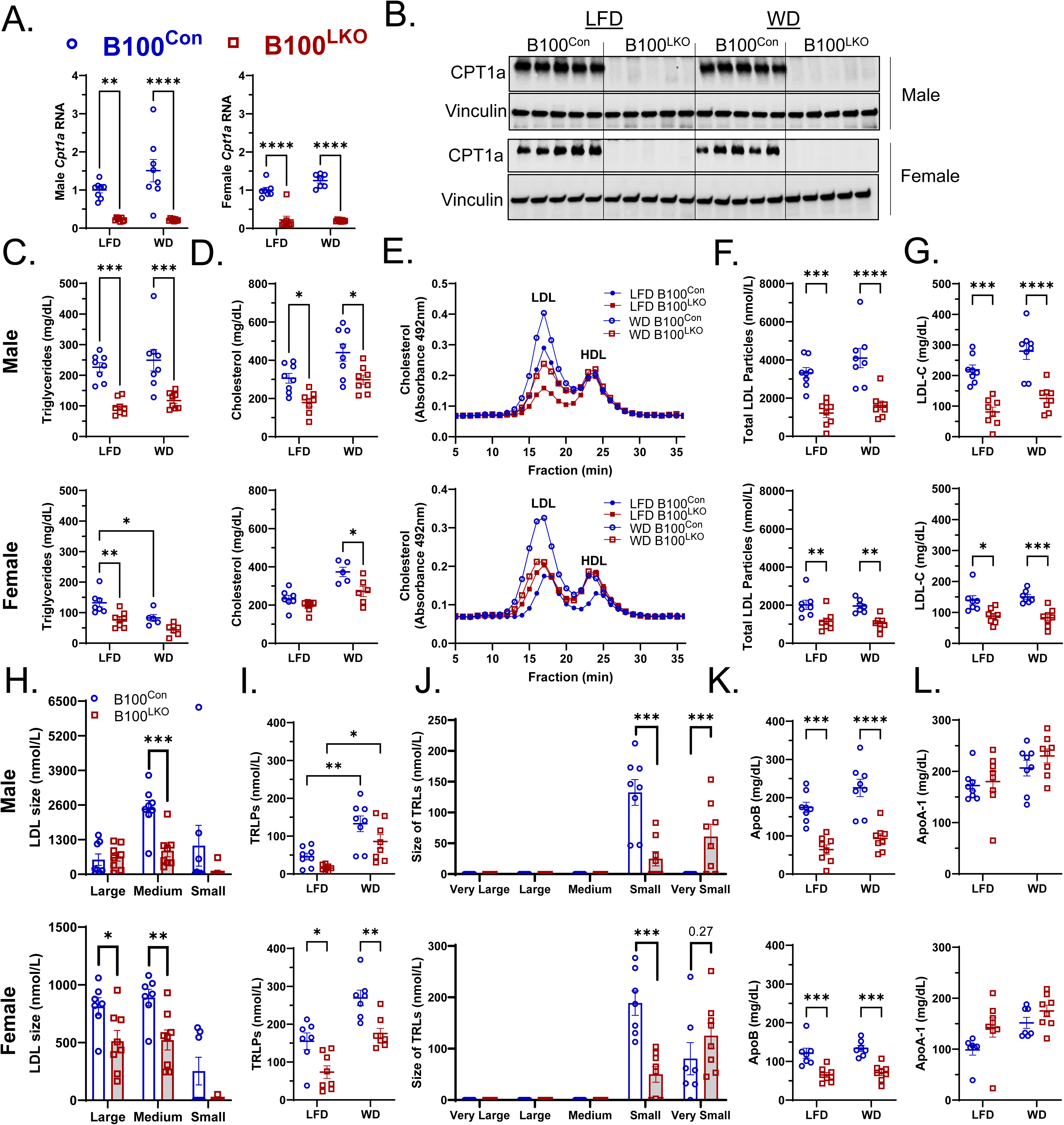
Liver-specific Cpt1a Deletion Reduces LDL-C, LDL particle number, and ApoB Concentrations in Human ApoB100-transgenic Mice. Male and female B100·*Cpt1a^F/F^* mice were treated with AAVs containing empty (B100^Con^) or TBG-Cre (B100^LKO^) prior to WD-feeding for 16-weeks. Mice were necropsied in the fed state. (**A-B**) Liver Cpt1a RNA and protein levels were measured by qPCR (**A**; n=7-8) and western blot (**B**; n=5), respectively. Vinculin serves as a loading control. (**C-G**) Steady-state triglycerides and cholesterol in whole plasma (**C**, **D**) and cholesterol levels in FPLC-fractionated plasma (**E**) from B100^Con^ and B100^LKO^ mice. (**F-L**) NMR analysis of B100^Con^ and B100^LKO^ plasma for total LDL particle concentration (**F**), LDL-C (**G**), LDL particle size (**H**), TRLPs (**I**), TRLP size (**J**), ApoB (**K**), and ApoAI (**L**). Data are presented with males on top and females on bottom. Significance was determined either by a two-way ANOVA with Tukey’s multiple comparison *post hoc* analysis or a Student’s two-tailed *t*-test as shown in panels H and J. *P<0.05; **P<0.01; ***P<0.001; ****P<0.0001.

We then examined plasma cholesterol levels and in response to WD feeding, B100^Con^ mice had increased plasma cholesterol levels in both male and female mice (diet [male] *p*=0.0002; diet [female] *p*<0.0001; **Fig. 4D, E**). This effect is almost exclusively in apoB-LPs (**Fig. 4E**). Inactivation of Cpt1a reduced total plasma cholesterol levels in both sexes irrespective of diet (genotype [male] *p*=0.0001; genotype [female] *p*=0.0018; **Fig. 4D**). These reductions were limited to apoB-LPs, as no differences in HDL-C was observed by size exclusion chromatography (**Fig. 4E**). Consistently, male and female B100^LKO^ mice exhibited ∼40-60% reductions in total LDL particle number (**Fig. 4F**) and LDL-C (**Fig. 4G**) as measured by NMR.

Several reports have indicated that small dense LDL (sdLDL) particles are more greatly associated with atherogenesis and metabolic disorders across independent human cohorts, as compared to total LDL particle concentrations (70–72). Therefore, we examined the size distribution of LDL particles in the circulation of WD-fed B100^Con^ and B100^LKO^ mice. Both male and female B100^LKO^ mice exhibited significant reductions in medium-sized LDL particles (20.5-21.4 nm; **Fig. 4H**) while small LDL particles (19-20.4 nm) trended to decrease in LKO mice irrespective of sex (**Fig. 4H**). Female B100^LKO^ mice also exhibited significant reductions in large LDL (21.5-23 nm) while these particles were unchanged in male B100^LKO^ mice (**Fig. 4H**).

Triglyceride rich lipoproteins (TRLPs) are predominately composed of triglycerides and arise from the liver as VLDL or intestine as chylomicrons. These secreted TRLPs are catabolized by lipases in the vasculature leading to smaller apoB-LPs. Similar to total plasma cholesterol levels in **Fig. 4D**, WD-feeding increased TRLP levels in both male and female B100^Con^ mice. However, liver-specific deletion of Cpt1a reduced TRLP concentrations by 40-50% in both sexes (genotype [male] *p*=0.013; [female] *p*<0.0001) irrespective of diet (**Fig. 4I**). Male and female B100^LKO^ mice exhibited significant reductions in small TRLPs (30-36 nm) with increases in very small TRLPs (24-29 nm; **Fig. 4J**), supporting the notion of accelerated metabolism of particles. No very large (90-240 nm), large (50-89 nm), or medium (37-49 nm) sized TRLPs were detected in male or female mice. Consistent with reductions in both TRLPs and LDL-C, steady-state apoB concentrations decreased by ∼50% independent of sex and diet in B100^LKO^ mice (**Fig. 4K**), while no differences in ApoA1 were observed between genotypes (**Fig. 4L**).

### B100^LKO^ Mice Exhibit No Changes in Biliary Cholesterol Secretion Rates, Bile Acid Concentrations, or Fecal Sterol Loss

To determine if reductions in plasma cholesterol were associated with accelerated sterol elimination, we examined biliary secretion of neutral and acidic sterols and fecal neutral sterols. At necropsy, the gall bladder was cannulated and bile diverted bile into collection tubes for a period of 30 minutes to asses basal bile flow and composition (43). As expected, WD feeding increased biliary cholesterol secretion, but bile flow, cholesterol secretion, and bile acid secretion rates were unaffected by genotype (**Fig. 5A-C**). The concentration of total bile acids in the gall bladder were unchanged in both male and female B100^LKO^ mice (**Fig. 5D**). We also analyzed bile acid composition. Male B100^LKO^ mice had significant reductions in deoxycholic acid while females had modest, but significant reductions in muricholic and ursodeoxycholic acid (**Fig. 5E**). Despite these differences, neutral sterol loss into the feces was unaffected by genotype (**Fig. 5F**). Taken together, steady-state reductions in cholesterol levels observed in B100^LKO^ mice cannot be attributed to accelerated sterol secretion into bile or excretion into the feces.

**Figure 5.**
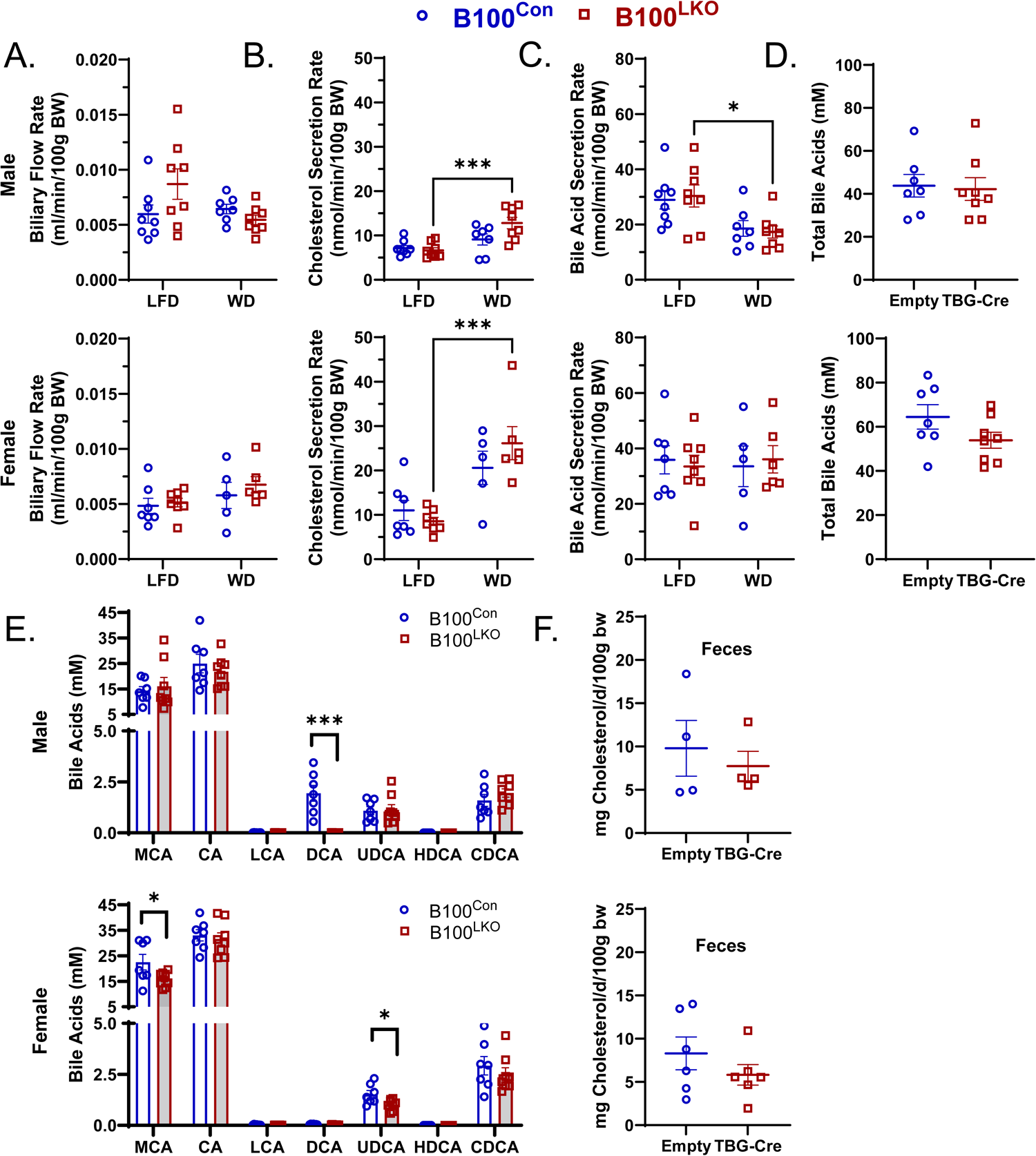
Sterol Excretion is Unaffected by Hepatic Cpt1a Deletion. Male and female B100^Con^ and B100^LKO^ mice were fed a WD for 16-weeks. (**A-C**) Bile flow (**A**), cholesterol secretion (**B**), and bile acid secretion (**C**) rates were determined over a 30-minute time period (n=7-8). (**D-E**) Gall bladder bile acid concentration (**D**) and composition (**E**) were determined by UPLC-MS/MS (n=7-8). (**F**) Fecal cholesterol was measured by GC-MS. Significance was determined either by a two-way ANOVA with Tukey’s multiple comparison *post hoc* analysis or a Student’s two-tailed *t*-test as shown in panel E. Data are presented with males on top and females on bottom. *P<0.05; ***P<0.001.

### VLDL-TG and VLDL-C Secretion Rates Are Accelerated in Female B100^LKO^ Mice

Reductions in steady-state levels of plasma lipoproteins can be driven by two primary mechanisms, reduced secretion or accelerated clearance from the plasma compartment. We first measured VLDL-TG and VLDL-C secretion rates following inhibition of LPL with poloxamer 407 (39). Despite reduced steady-state levels of apoB-LPs, VLDL-TG secretion was unaffected or slightly elevated in male B100^LKO^ mice (*p*=0.324; **Fig. 6A**) with no difference in VLDL-C secretion rate (**Supplemental Figure 6A**). On the contrary, female B100^LKO^ mice had accelerated VLDL-TG (*p*=0.024; **Fig. 6B**) and tended towards accelerated VLDL-C secretion (*P*=0.081; **Supplemental Figure 6B**) compared to female B100^Con^ mice. Despite greater VLDL secretion rates, female B100^LKO^ mice had lower human apoB100 concentrations (via ELISA) 6-hours after LPL inhibition indicating female B100^LKO^ mice are secreting fewer, but larger and more lipidated particles (**inset to Fig. 6A, B**).

**Figure 6.**
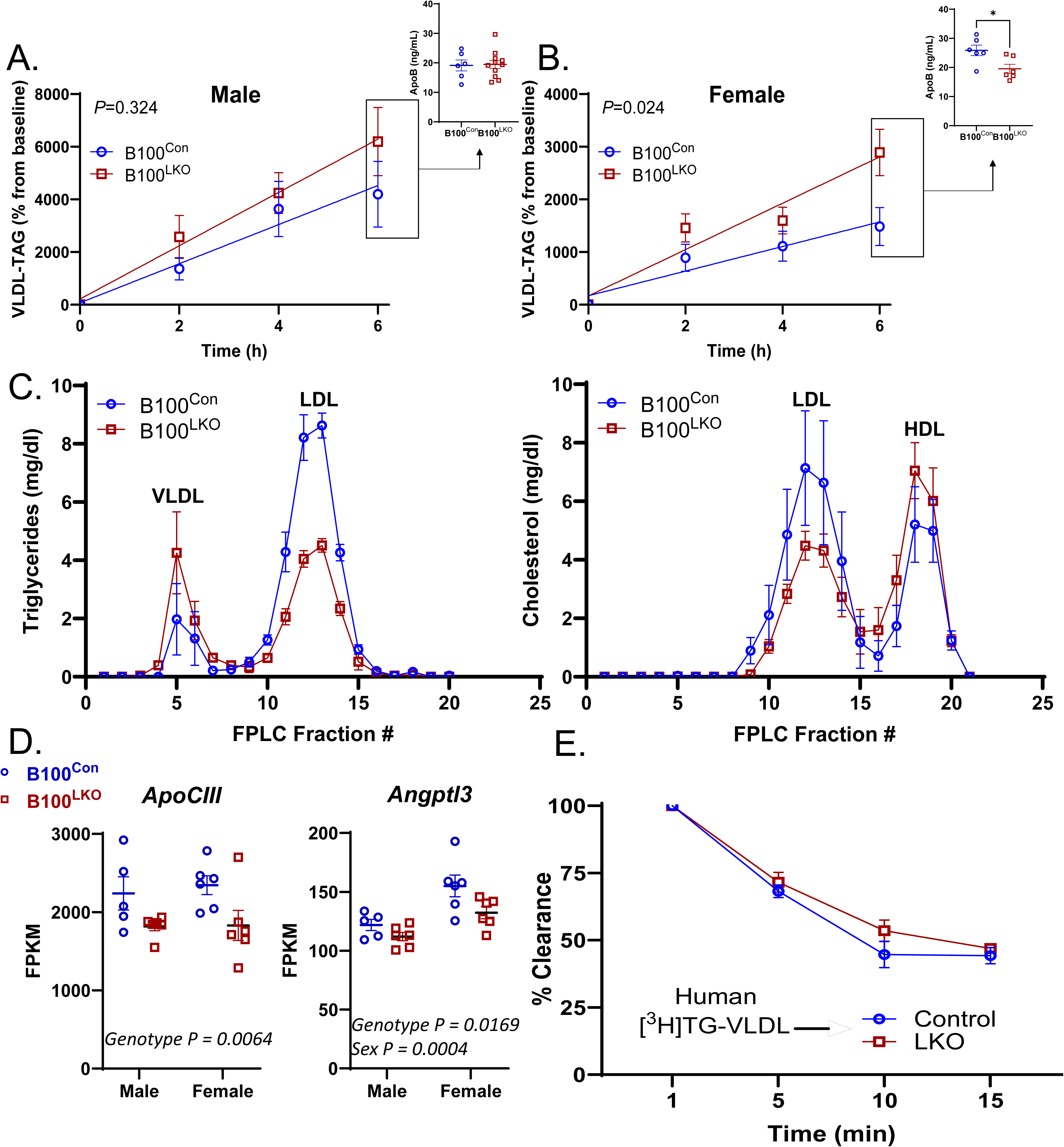
VLDL-TG Secretion and Metabolism is Accelerated in B100^LKO^ Mice. (**A, B**) Plasma triglycerides were measured in male (**A**) and female (**B**) B100^Con^ and B100^LKO^ mice after P407 treatment (*n*=6-11). Human apoB100 was measured via ELISA using plasma collected from mice 6-hours post P407 treatment (inset; *n*=6-11). (**C**) Blood was collected from the interior vena cava between the hepatic vein and right atrium of male B100^Con^ and B100^LKO^ mice. Triglycerides and cholesterol were then measured in FPLC-fractionated plasma (*n*=3). (**D**) Liver bulk RNA-sequencing reads for ApoCIII and Angptl3 from male and female B100^Con^ and B100^LKO^ mice (*n*=5-6). (**E**) Male B100^Con^ and B100^LKO^ mice were injected with human [^3^H]-triolein labeled VLDL and blood sampled at 1, 5, 10, and 15 minutes was used for scintillation counting. Data are expressed as percent initial counts at 1 minute (*n*=5-6). Significance was determined either by a two-way ANOVA with Tukey’s multiple comparison *post hoc* analysis (panel E) or a Student’s two-tailed *t*-test as shown in panel B (inset). *P<0.05

The reductions in steady-state triglyceride and cholesterol levels in LKO mice (**Figs. 2, 4**) were observed in plasma isolated via cardiac puncture. The rapid metabolism of TRLPs is such that newly synthesized lipoproteins emerging from the liver are not typically observed in plasma collected from peripheral blood. For this reason, we collected plasma from the inferior vena cava between the hepatic vein and right atrium in an independent cohort of fasted male B100^Con^ and B100^LKO^ mice fed a WD for 8-weeks. From this location, we can observe the composition of VLDL before (fractions 4-6) and after (fractions 10-14) peripheral metabolism. Consistent with LPL inhibitor studies in **Fig. 6A**, male B100^LKO^ mice had slightly elevated VLDL-TGs (**Fig. 6C**). However, LDL-TGs and LDL-C were substantially reduced in the fractions after peripheral metabolism (fractions 10-14; **Fig. 6C**). Consistent with mice lacking the human B100 transgene (**Fig. 3**), normalized sequencing reads revealed significant reductions in the expression of both ApoCIII and Angptl3 in B100^LKO^ compared to B100^Con^ mice, two well-established inhibitors of LPL-mediated metabolism of apoB-LPs (**Fig. 6D**).

To determine if reductions in steady-state apoB-LPs are due to a general increase in the peripheral metabolism of TG-rich apoB-LPs, we radiolabeled human VLDL with [^3^H]-triolein and analyzed clearance rates in B100^Con^ and B100^LKO^ mice. Plasma clearance of human VLDL-TG was not affected by Cpt1a deletion (**Fig. 6E**), suggesting that the factors intrinsic to endogenously-produced VLDL accelerate apoB-LP clearance in LKO and B100^LKO^ mice.

### Female B100^LKO^ Mice Exhibit Free Cholesterol Accumulation, Inflammation, and Fibrosis of the Liver

Given the significant reductions in circulating apoB-LPs in response to hepatic CPT1a-deletion (**Figs. 2, 4**), we next determined whether hepatic lipid levels would also be affected. Overall liver weights and liver weight-to-body weight ratios were increased in both WD-fed male B100^Con^ and B100^LKO^ as compared to LFD-fed mice (**Supplemental Figure 7A**). Consistently, male B100^Con^ and B100^LKO^ accumulated significantly (*p*<0.0001) greater triglycerides, total and esterified cholesterol in response to WD-feeding as compared to LFD-fed mice (**Fig. 7A-D**). However, similar to **Figure 2I**, no differences were observed between genotypes in male mice maintained on either diet. Conversely, liver weight is elevated in female B100^LKO^ mice regardless of diet (**Supplemental Figure 7B**). Hepatic triglycerides increased two-fold from 124.5 μg/mg to 265.9 μg/mg of liver weight as compared to WD-fed B100^Con^ mice (**Fig. 7A**). Female WD-fed B100^LKO^ mice also accumulated 2-2.5 fold greater total, esterified, and free cholesterol levels in the liver as compared to controls (**Fig. 7B-D**). Biochemical and histological analysis revealed that female control mice are protected from steatosis compared to their male counterparts (**Fig. 7A, E**). Inactivation of Cpt1a sensitized female mice to WD feeding-induced steatosis characterized by the accumulation of small lipid droplets surrounding the central vein (**Fig. 7E**).

**Figure 7.**
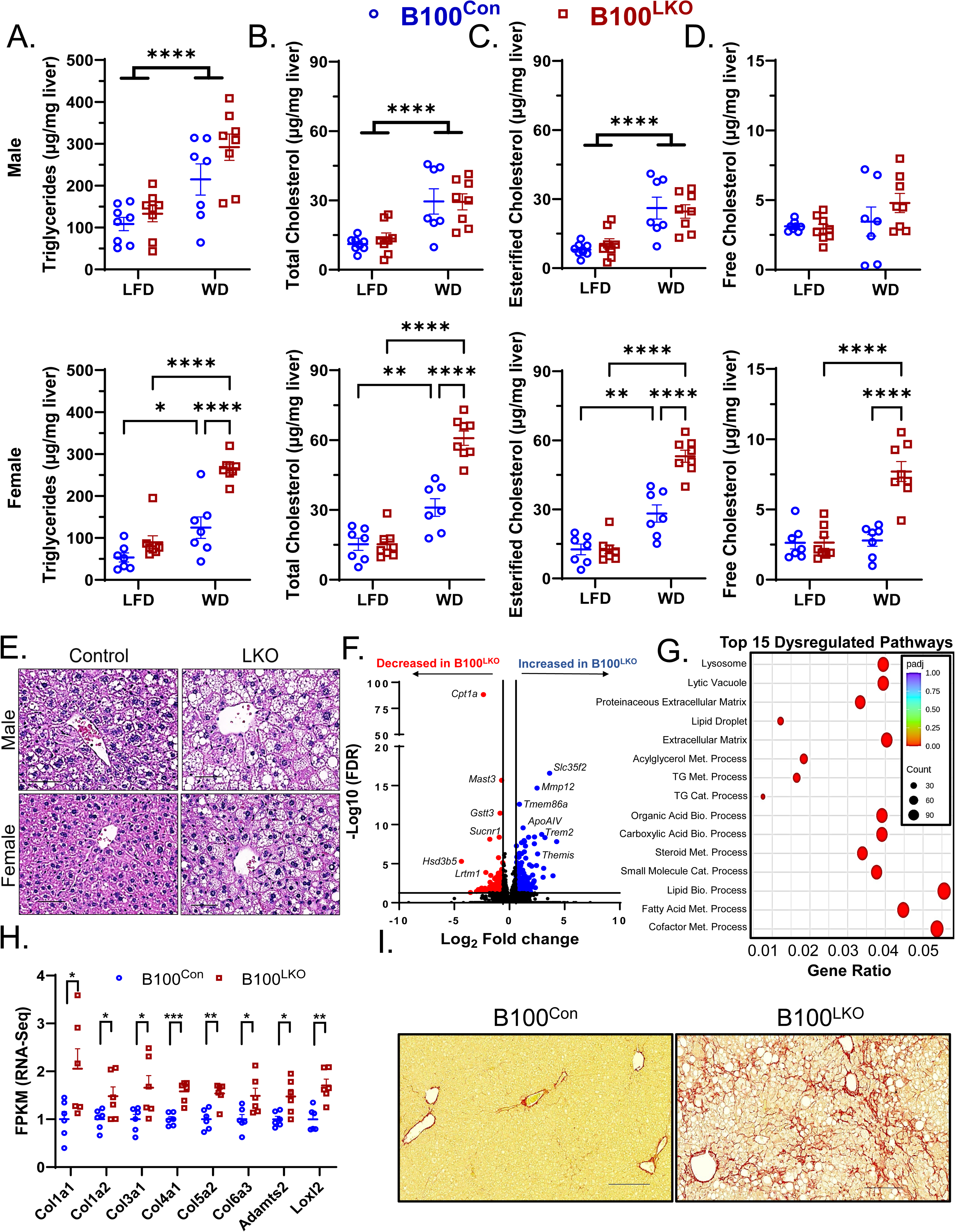
Female B100^LKO^ Mice Accumulate Free Cholesterol and Exhibit Fibrosis of the Liver. Male and female B100^Con^ and B100^LKO^ mice were fed a WD for 16-weeks. (**A-D**) Hepatic triglycerides (**A**), total cholesterol (**B**), esterified (**C**) and free (**D**) cholesterol levels were measured (*n*=7-8). (**E**) H&E staining of livers from male and female B100^Con^ and B100^LKO^ mice. The central vein is shown in each image. (**F**) A volcano plot showing differential gene expression comparing female B100^Con^ and B100^LKO^ mice fed a WD. Genes increased (in blue) or decreased (in red) with Cpt1a-deficiency. The horizontal black bar denotes the significance cutoff FDR=0.05. The vertical black bars denote a minimal threshold for the effect size of 1.5 (Log2 fold change=±0.58). (**G**) KEGG analysis dot plots for the top 15 dysregulated pathways in female B100^Con^ verse B100^LKO^ mice. (**H**) Total read counts (in FPKM) for genes involved in collagen deposition in the liver (*Col1a1, Col1a2, Col3a1, Col4a1, Col5a2, Col6a3, Adamts2, Loxl2*; *n*=6). (**I**) Picrosirius red staining on liver tissue from female B100^Con^ and B100^LKO^ mice. Significance was determined by a two-way ANOVA with Tukey’s multiple comparison *post hoc* analysis (panels A-D) or a Student’s two-tailed *t*-test as shown in panel H. *P<0.05; **P<0.01; ***P<0.001; ****P<0.0001.

Free cholesterol accumulation in hepatocytes can exacerbate diet-induced metabolic dysfunction-associated steatohepatitis (MASH) and fibrosis (73). Therefore, we completed bulk RNA-sequencing on whole livers from WD-fed female mice to determine whether the free cholesterol accumulation observed in female WD-fed B100^LKO^ mice (**Fig. 7D**) associates with greater inflammation. We identified 306 genes that met pre-determined statistical (p-adjusted value ≤0.05) and effect size cut-offs (Log2 normalized fold-change [FC] of ±0.58). Of the 306 genes identified, 193 were upregulated while 113 were downregulated in response to *Cpt1a* deficiency (**Fig. 7F**). The most downregulated gene in female B100^LKO^ mice was Cpt1a, while several other genes involved in succinate (*Sucnr1*), and xenobiotic and steroid metabolism (*Gstt3, Hsd3b5*) were significantly downregulated compared to control mice (**Fig. 7F**). The most significantly upregulated gene in female B100^LKO^ mice was the solute carrier family 35 member F2 (*Slc35f2*) followed by genes associated with T-lymphocyte development (*Themis*) and the accumulation of lipid-loaded macrophages (*Trem2, Tmem86a*; **Fig. 7F**) (74, 75).

To determine pathways differentially affected by *Cpt1a*-deletion in the liver, we completed a KEGG enrichment analysis. Out of the top 15 dysregulated pathways comparing female B100^Con^ and B100^LKO^ mice, 9 were related to steroid and fatty acid metabolism. Notably, 2 pathways significantly enriched in B100^LKO^ mice were related to extracellular matrix remodeling (**Fig. 7G**). We then analyzed normalized read counts for individual genes involved in extracellular matrix remodeling pathways, and observed significant increases in collagen genes (*Col1a1, Col1a2, Col3a1, Col4a1, Col5a2, Col6a3*), and genes involved in collagen biosynthesis (*Adamts2*) and crosslinking (*Loxl2*) in livers from female WD-fed B100^LKO^ mice (**Fig. 7H**). Consistent with changes at the gene expression level, picrosirius red staining revealed significant fibrosis in livers from female B100^LKO^ mice (**Fig. 7I**). Taken together, female WD-fed B100^LKO^ mice accumulate 2.5X greater free cholesterol levels which associate with an exacerbation of inflammation and fibrosis of the liver as compared to female B100^Con^ mice.

### Distinct B- and T-lymphocyte Populations Are Enriched in Livers from WD-fed Female B100^LKO^ Mice

Given the changes in inflammatory gene expression by bulk RNA-sequencing (**Fig. 7F-H**), we went on to perform single cell RNA-sequencing on parenchymal and non-parenchymal cell populations from WD-fed female B100^Con^ and B100^LKO^ mice. Uniform Manifold Approximation and Projection (UMAP) techniques showed distinct clustering of T-lymphocytes, natural killer (NK) cells, B-lymphocytes, Kupffer Cells, Macrophages, Fibroblasts, Endothelial Cells, and Hepatocyte cell populations in the liver (**Fig. 8A**). This clustering was based off common gene expression patterns across cell populations as shown in **Supplemental Figure 8**. Previous studies have shown stepwise reductions in hepatocyte populations throughout the progression from MASLD to advanced MASH (55). Consistently, a direct UMAP comparison across genotypes revealed a 50% reduction in the total hepatocyte population in B100^LKO^ mice (**Fig. 8B-D**). Female B100^LKO^ mice also exhibited a 40% reduction in intrahepatic T-lymphocyte population 1, an anti-tumor effector-like CD8+ T cell population characterized by enrichment of the chemokine receptor *Cxcr6* (76) and immune receptor *Cd226* (77, 78) (**Fig. 8B-D**). On the contrary, female B100^LKO^ mice accumulated 30% more B lymphocytes and ∼3X more T lymphocyte population 5, which is enriched in the proinflammatory chemokine *Ccl5* (otherwise known as RANTES) and in the cytotoxic T cell marker *Cd8b1* (**Fig. 8B-D**).

**Figure 8.**
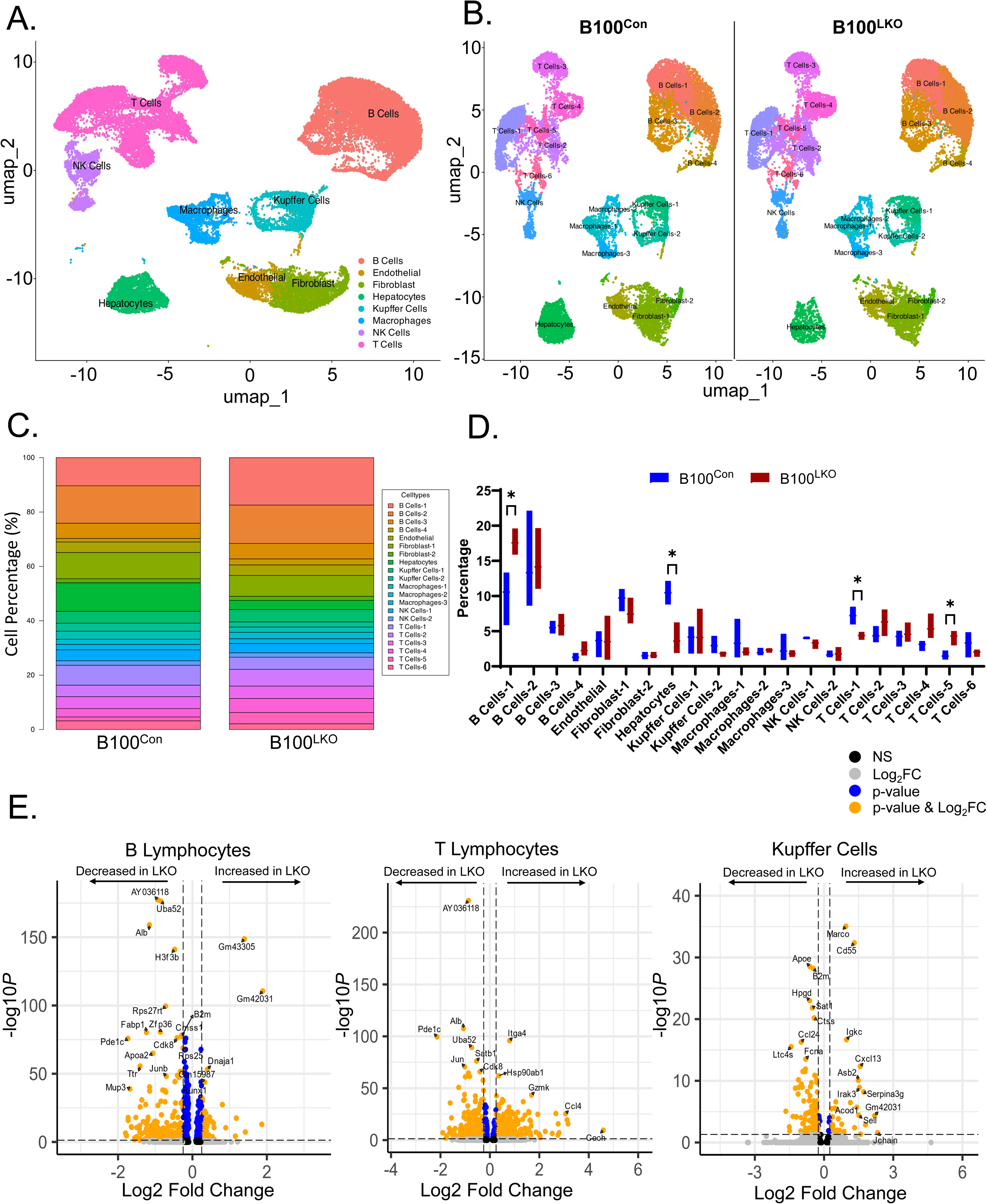
Distinct B- and T-lymphocyte Populations Accumulate in Livers of Female B100^LKO^ Mice. Single cell RNA-sequencing profiling of parenchymal and nonparenchymal cells of female B100^Con^ and B100^LKO^ mice fed a WD for 22 weeks. (**A**) UMAP visualizations of mouse liver cells isolated from both B100^Con^ and B100^LKO^ mice, and clustered across 8 cell populations (*n*=6 total). (**B**) UMAP visualizations of liver cells clustered across 21 defined cell populations comparing B100^Con^ verse B100^LKO^ mice (*n*=3/genotype). (**C, D**) The overall percentage (**C**) of each of the 21 defined cell types (**D**) in the livers from female B100^Con^ verse B100^LKO^ mice. (**E**) A volcano plot showing differential gene expression across different cell populations (B-Lymphocytes, T-Lymphocytes, Kupffer Cells) comparing female B100^Con^ and B100^LKO^ mice fed a WD. Genes increased (on right) or decreased (on left) with *Cpt1a*-deficiency. The horizontal black bar denotes the significance cutoff FDR=0.05. Significance was determined by a Student’s two-tailed *t*-test as shown in panel D. *P<0.05.

We then went on to identify all DEGs across B- and T-lymphocytes, and Kupffer cell populations in livers from WD-fed female B100^Con^ and B100^LKO^ mice (**Fig. 8E**). In B-lymphocytes, 134 genes met pre-determined statistical (p-adjusted value ≤0.05) and effect size cut-offs (Log2 normalized fold-change [FC] of ±0.58). Of the 134, only 9 genes were increased in livers of B100^LKO^ mice while 252 were elevated in B100^Con^ mice (**Fig. 8E**). Several of the notable genes decreased in the B-cell population from B100^LKO^ mice were *Junb*, a negative regulator of B-cell proliferation and transformation (79), and *Zfp36*, a regulator of quiescence in developing B-lymphocytes (80). In T-lymphocytes, 261 genes met pre-determined statistical (p-adjusted value ≤0.05) and effect size cut-offs (Log2 normalized fold-change [FC] of ±0.58). Of the 261 genes identified, 136 were increased while 125 were decreased in T-lymphocytes from livers of B100^LKO^ mice (**Fig. 8E**). Interestingly, female B100^LKO^ mice exhibited increased expression of granzyme K (*Gzmk*), a marker of age-associated inflammation (81), and *Ccl4* (otherwise known as macrophage inflammatory protein-1β), a proinflammatory chemoattractant for several immune cell types including T-cells, NK-cells, monocytes, and pro-tumorigenic macrophages (**Fig. 8E**) (82, 83). In Kupffer cells, 63 genes met pre-determined statistical (p-adjusted value ≤0.05) and effect size cut-offs (Log2 normalized fold-change [FC] of ±0.58). Of the 63 genes identified, 22 were increased while 41 were decreased in Kupffer cells from livers of B100^LKO^ mice (**Fig. 8E**). Notable genes increased in Kupffer cells of female B100^LKO^ mice were *Cxcl13*, a Kupffer cell proinflammatory chemokine that promotes accumulation of CD4+ T and B-cells (84), and *Marco*, a scavenger receptor class A protein expressed in more proinflammatory M1-like macrophages (**Fig. 8E**) (85). Taken together, female B100^LKO^ mice exhibit a more profound proinflammatory gene signature that associates with greater accumulation of B- and T-cell lymphocyte populations in the liver when challenged with a WD.

## DISCUSSION

There is empirical evidence that CPT1a expression and activity positively associates with plasma cholesterol levels across independent human cohorts (26, 57–59). For instance, greater methylation at two independent sites in *CPT1a* (cg00574958, cg17058475) associate with reductions in total VLDL and LDL particle number (59). On the contrary, FH mutation-negative patients (e.g. FH patients who are non-carriers of mutations in *LDLR*, *APOB*, or *PCSK9*) exhibit *CPT1a* hypomethylation (cg00574958) while no differences in methylation of other lipid-modifying genes were identified (26).

The major focus of this work was to provide mechanistic insight explaining these associations. First, we show via the Common Metabolic Diseases Knowledge Portal that the *Cpt1a* missense variant Lys455Thr (rs189174414) was strongly inversely associated with both total cholesterol (β = −0.6) and LDL-C (β = −0.2), and has been shown to result in impaired folding and degradation of the protein (86). Other loss-of-function mutations, such as the missense variant Ala275Thr (rs2229738) (60), were also negatively associated with plasma cholesterol and LDL-C levels. Consistent with human associations, data from the HMDP demonstrates significant positive associations with hepatic Cpt1a RNA levels and total cholesterol and LDL-C levels across inbred strains of mice. These results implicate CPT1a as a potential modulator of plasma cholesterol in humans and mice.

The liver is the primary source of ketone bodies in the fasted state, a process that is reliant on the generation of acetyl-CoA via FAO. Not surprisingly, we observed significant reductions in plasma β-hydroxybutyrate in fasted LKO mice. Reduced availability of acyl-CoA is known to reduce the acetylation of proteins, including histones (87). In fact, lysine acetylation of mitochondrial proteins is largely driven by acetyl-CoA derived from mitochondrial FAO of acyl-CoAs (88). On the contrary, Cpt1a deletion in the liver could result in accumulation of acyl-CoAs which could further alter protein acylation. The role of protein acylation as a mediator of cellular responses to metabolic perturbations, including enzymatic activity, protein stability, and transcription is an emerging area of research [reviewed in (89)]. Cpt1a is also reported to have succinyltransferase activity (90), and reduced flux of acyl-CoA into the TCA cycle would be expected to reduce the availability of TCA intermediates including succinate. The impact of Cpt1a on the hepatic acetylome and succinylome have yet to be investigated, but could be additional mechanisms by which FAO-derived substrates further modulate lipid and apoB-LP metabolism. By example, acetylation of sterol regulatory binding protein (SREBP)-1 and SREBP2, master regulators of lipogenesis and cholesterol synthesis respectively, regulate their stability and transcriptional activity (91). A proteomic analysis of the acetylome and succinylome from human embryonic stem cells differentiated to a hepatocyte-like phenotype revealed acetylation of multiple apolipoproteins (AI, B, E, CIII, AIV) (92), but neither the impact of Cpt1a on the acetylation of apolipoproteins nor the effect of acetylation on lipoprotein assembly, secretion or clearance is known.

The transcriptional regulator PPARα mediates the adaptive response to fasting. Our KEGG analysis indicated enrichment of PPARα regulated genes in both the fasted and fed state in mice lacking hepatic Cpt1a. This is consistent with increased nuclear PPARα regardless of nutritional status and an enrichment of PPARα motifs in our H3K27ac ChIP-seq data set. What is driving enhanced PPAR signaling in the absence of Cpt1a is not yet known. One possibility is increased abundance of FFAs, which act as PPAR ligands, in cells that lack the ability to efficiently oxidize them in mitochondria. Alternatively, reduced mitochondrial β-oxidation in the absence of Cpt1a in both the fasted and fed states may generate signals that promote PPAR signaling in a cellular effort to restore oxidative metabolism. Consistent with this idea, the alternative FAO pathway in peroxisomes is elevated in the absence of Cpt1a likely due to increased PPARα signaling. In a type 1 diabetic mouse model, hepatic peroxisomal FAO generates free acetate which is used as a precursor for *de novo* cholesterol synthesis (93). However, the contribution of peroxisomal FAO to apoB-LP production and clearance is currently unknown and warrants further investigation.

The reduction of apoB-LPs in Cpt1a LKO mice was not associated with a reduction in VLDL secretion, but rather a modest increase. In females, this increase was associated with reduced apoB levels in plasma, indicating the secretion of larger, TG-rich VLDLs. This appears to be mediated in part by an increase in levels of ApoAIV which is known to promote the expansion of VLDL particles secreted by the liver (94). Unlike the previous study, the increase in ApoAIV and accelerated VLDL secretion was not associated with a reduction in hepatic TGs, particularly in female mice. A recent report indicates zonal and sexually dimorphic expression of the VLDL receptor in the liver with expression limited to central areas and greater expression in females compared to males (95). Conversely, VLDL assembly was more prominent in portal areas, suggesting portal secretion and central reuptake of VLDL in the liver, particularly in female mice. In the present study and in our previously reported work (15), the absence of Cpt1a promoted hepatic TG and cholesterol accumulation to a much greater extent in females. This could potentially be explained by accelerated secretion and reuptake of VLDL. However, hepatic Cpt1a deficiency promoted pan-lobular microsteatosis in female mice (15). Zonal expression of Cpt1a, the mechanism(s) by which Cpt1a influences lipid droplet size in female mice, and the impact of zonal-Cpt1a deficiency on hepatic lipid and lipoprotein metabolism awaits further investigation.

Our data shows that LKO mice exhibit greater clearance of endogenously-produced VLDL particles, as opposed to a general increase in LPL-mediated clearance of apoB-LPs. This is supported by two lines of evidence: 1) plasma collected from the inferior vena cava between the hepatic vein and right atrium show an increase in newly synthesized VLDL-TGs with reductions in LDL-TGs and LDL-C, and 2) exogenous VLDL exhibited similar clearance rates between control and LKO mice. Factors such as size and composition of endogenously-produced VLDL particles, and their remnants, may further affect their clearance rates. For example, we previously demonstrated that Cpt1a deficiency in the liver alters the phospholipid pool (PC/PE ratios) by decreasing expression of phosphatidylethanolamine N-methyltransferase (*Pemt*) (15). Mice with Pemt deficiency exhibit significant reductions in PC enrichment into nascent VLDL particles, which coincides with greater VLDL diameter and accelerated clearance of these particles (96). Consistently, our studies suggest that fewer, yet more lipidated VLDL particles are produced in response to Cpt1a deletion. This is supported by a ∼2.5X increase in VLDL-TG secretion rates while apoB levels were reduced 6-hours post-LPL inhibition in LKO mice. Therefore, it is plausible that Cpt1a deficiency alters the nascent VLDL phospholipid pool to facilitate changes in apolipoprotein binding and accelerated clearance of apoB-LPs. Current studies are underway to formally test these hypotheses.

While the data presented herein are supported by mouse models and human association data, the concept that reducing long chain FAO in the liver leads to reductions in plasma lipids (cholesterol, TGs) is quite intriguing, and in some ways, counterintuitive. However, our work herein aligns with that of independent groups showing increases in mitochondrial FAO via ACC inhibition (ACCi) leads to elevated plasma TGs and LDL-apoB fractional replacement rates (3–11). The increase in plasma TGs with ACCi is attributed to increases in both VLDL-TG secretion and impairments in VLDL clearance, consistent with elevations in circulating ApoCIII and Angptl3 (6). Consistent with our data, the phenotypes observed with ACCi can be rescued by PPARα agonist treatment (5, 6). Supporting evidence from this work shows that loss of CPT1a-mediated FAO leads to accelerated VLDL-TG and VLDL-C secretion rates, lower steady-state LDL-C and apoB concentrations, greater peripheral clearance of apoB-LPs, activation of hepatic PPARα, and decreases in ApoCIII and Angptl3. Mice deficient in ApoCIII exhibit normal VLDL secretion rates yet have accelerated clearance of both TG- and cholesteryl oleate-labeled particles (97). Further, loss-of-function mutations in *Angptl3* leads to reductions in plasma cholesterol levels which has been attributed to greater clearance of lipolytic remnants resulting in reductions in LDL-C (98). Taken together, reductions in hepatic ApoCIII and Angptl3 levels in response to reduced CPT1a-mediated FAO may explain greater apoB-LP clearance and lower steady-state plasma lipid levels in these mice.

Another intriguing phenotype observed herein and in our previous study (15) is the sex-specific accumulation of free cholesterol in livers of female LKO mice. This is of critical importance as free cholesterol accumulation promotes lipotoxicity-associated inflammation and exacerbation of liver fibrosis (73). These phenotypes are driven, in part, by a transcriptional regulator (TAZ) that induces expression of a host of pro-inflammatory and pro-fibrotic responses in response to free cholesterol accumulation (99). Thus, the free cholesterol accumulation observed in female LKO mice likely contributes to the exacerbation of inflammation and fibrosis in those animals. However, why deficiency of Cpt1a in hepatocytes results in free cholesterol accumulation in female but not male mice is currently unknown. Previous studies have noted that estrogens can increase FAO (100, 101) and decrease intracellular free cholesterol levels in human hepatoma cells (101, 102). Mechanistically, estrogens have also been shown to decrease acyl-coenzyme A:cholesterol acyltransferase 2 (ACAT2) activity in monkeys (103). Yet, the interrelationship between estrogens, CPT1a-mediated FAO, and free cholesterol levels warrants further investigation.

Taken together, these studies yield mechanistic insight linking loss-of-function variants in *CPT1a* to reductions in plasma cholesterol levels. Loss of Cpt1a protein levels in the livers of WT and human apoB100-transgenic mice lead to reductions in total plasma cholesterol, LDL-C, LDL particle number, and apoB levels. The reductions in steady-state cholesterol levels can be attributed to greater clearance of endogenously-produced VLDL particles and/or their respective remnants. Future studies should carefully consider the impact of Cpt1a-mediated FAO on cholesterol levels when considering new therapeutic options for the treatment of MASLD.

## DATA AVAILABILITY

All data are contained in the entirety of this manuscript. Additional information and requests for resources and reagents should be directed to and fulfilled by the Lead Contacts, Robert N. Helsley and Gregory Graf. Bulk RNA-sequencing data have been deposited in GEO under accession codes GSE277312 (Figure 3) and GSE277404 (Figures 6-7).

## ACKNOWLEDGEMENTS

We would like to acknowledge and thank the laboratories of Drs. Peter Carmeliet (Katholieke Universiteit Leuven) and Min Liu (University of Cincinnati) for sharing of the *Cpt1a*-floxed mice and ApoAIV antibody, respectively. This research was supported by services provided by the University of Kentucky Arts & Sciences Imaging Center. The UK pathology and light microscopy cores are supported in part by the Office of the Vice President for Research and the UK College of Medicine. We would also like to thank the University of Kentucky Center for Computational Sciences and Information Technology Services Research Computing for their support and use of the Morgan Compute Cluster and associated research computing resources.

## ABBREVIATIONS

CPT1a: Carnitine palmitoyltransferase 1a

LDL: low density lipoprotein

apoB-LPs: apoB-containing lipoproteins

PPARα: peroxisome proliferator activated receptor α

MASLD: metabolic dysfunction-associated steatotic liver disease

FH: familial hypercholesterolemia

LDL-C: LDL-cholesterol

ACC: acetyl-CoA carboxylase

FAO: fatty acid oxidation

EWAS: epigenetic wide association studies

GWAS: genome wide association studies

ChIP: chromatin immunoprecipitation

LDLR: low density lipoprotein receptor

LDLRAP: LDLR adaptor protein

PCSK9: proprotein convertase subtilisin/kexin type 9

apoB: apolipoprotein B

HDL-C: high density lipoprotein cholesterol

KEGG: Kyoto Encyclopedia of Genes and Genomes

MASH: metabolic dysfunction-associated steatohepatitis.

## TABLES WITH TITLES AND LEGENDS

**Supplemental Table 1.**
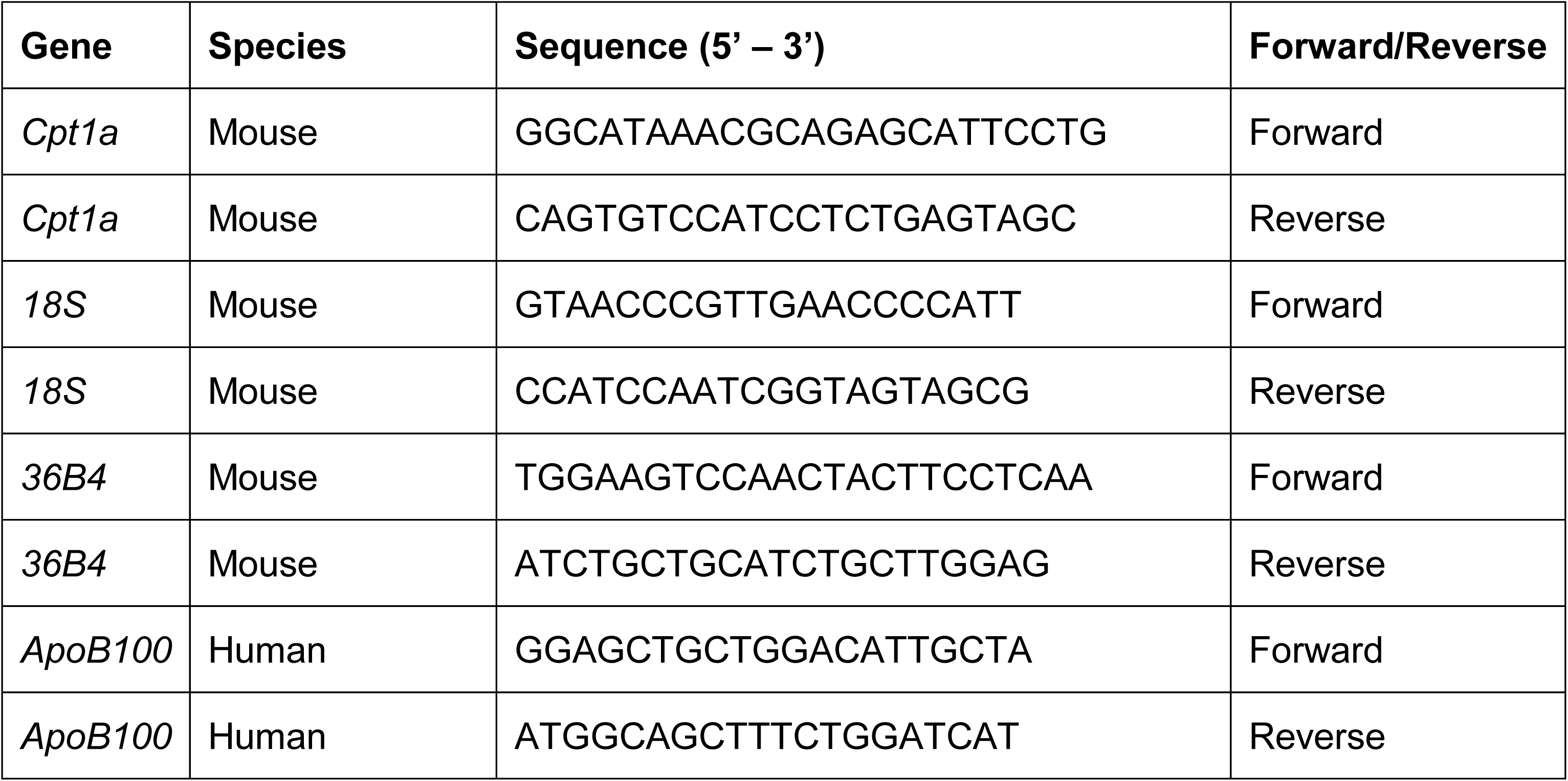
Primers for real-time PCR.

**Supplemental Table 2.** Antibodies for immunoblotting.

| **Protein** | **Catalog Information** | **Host Species & Clonality** | **Dilution** |
| --- | --- | --- | --- |
| CPT1a | 128568 | Mouse Monoclonal | 1:1000 |
| SREBP1 | NB100-2215 | Rabbit Polyclonal | 1:1000 |
| ChREBP | NB400-135 | Rabbit Polyclonal | 1:1000 |
| PPARα | sc-398394 | Mouse Monoclonal | 1:1000 |
| GAPDH | 5174 | Rabbit Monoclonal | 1:1000 |
| H3 | 4499 | Rabbit Monoclonal | 1:1000 |
| APOCIII | PA5-78802 | Rabbit Polyclonal | 1:1000 |
| ANGPTL3 | PA5-34750 | Mouse Polyclonal | 1:1000 |
| APOCII | PA5-102480 | Rabbit Polyclonal | 1:1000 |
| APOAIV | Gift from Min Liu (University of Cincinnati) | Goat | 1:1000 |
| CPT2 | ab71435 | Rabbit Polyclonal | 1:1000 |
| VLCAD | sc-271225 | Mouse Monoclonal | 1:1000 |
| LCAD | ab82853 | Rabbit Polyclonal | 1:1000 |
| SCAD | ab156571 | Rabbit Monoclonal | 1:1000 |
| PMP70 | ab3421 | Rabbit Polyclonal | 1:1000 |
| ACOX1 | sc-517306 | Mouse Monoclonal | 1:1000 |
| Vinculin | NB600-1293 | Mouse Monoclonal | 1:1000 |
| Actin | A5441 | Mouse Monoclonal | 1:1000 |

## SUPPLEMENTAL INFORMATION TITLES AND LEGENDS

**Supplemental Figure 1.**
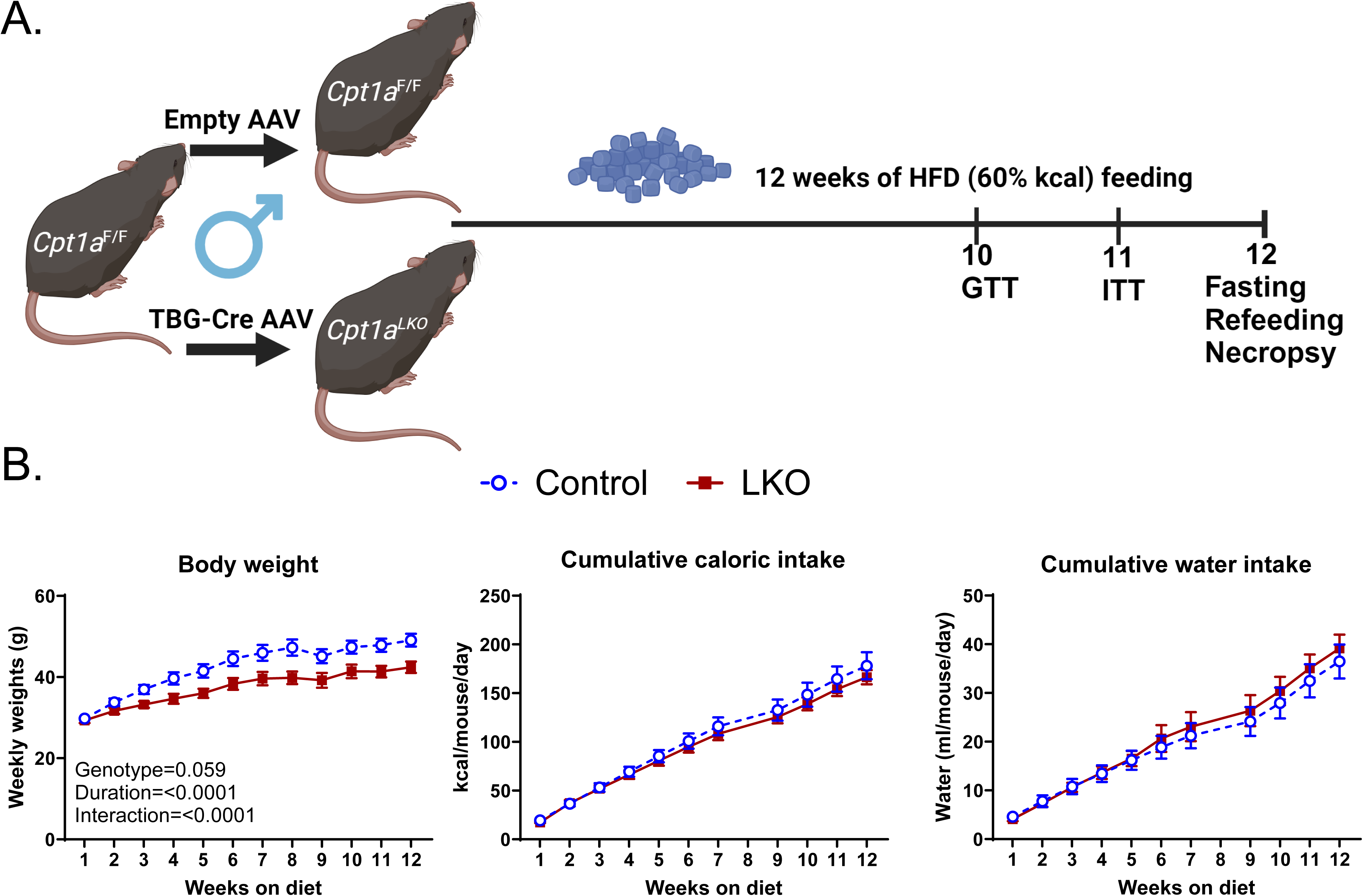
Liver-specific Deletion of Cpt1a Lowers Body Weight in Response to HFD-feeding. (**A**) An overview of the experimental design. (**B**) Body weight, cumulative caloric and water intake were measured throughout the experiment (*n*=6). Significance was determined by a repeated measures ANOVA. The ANOVA results are provided.

**Supplemental Figure 2.**
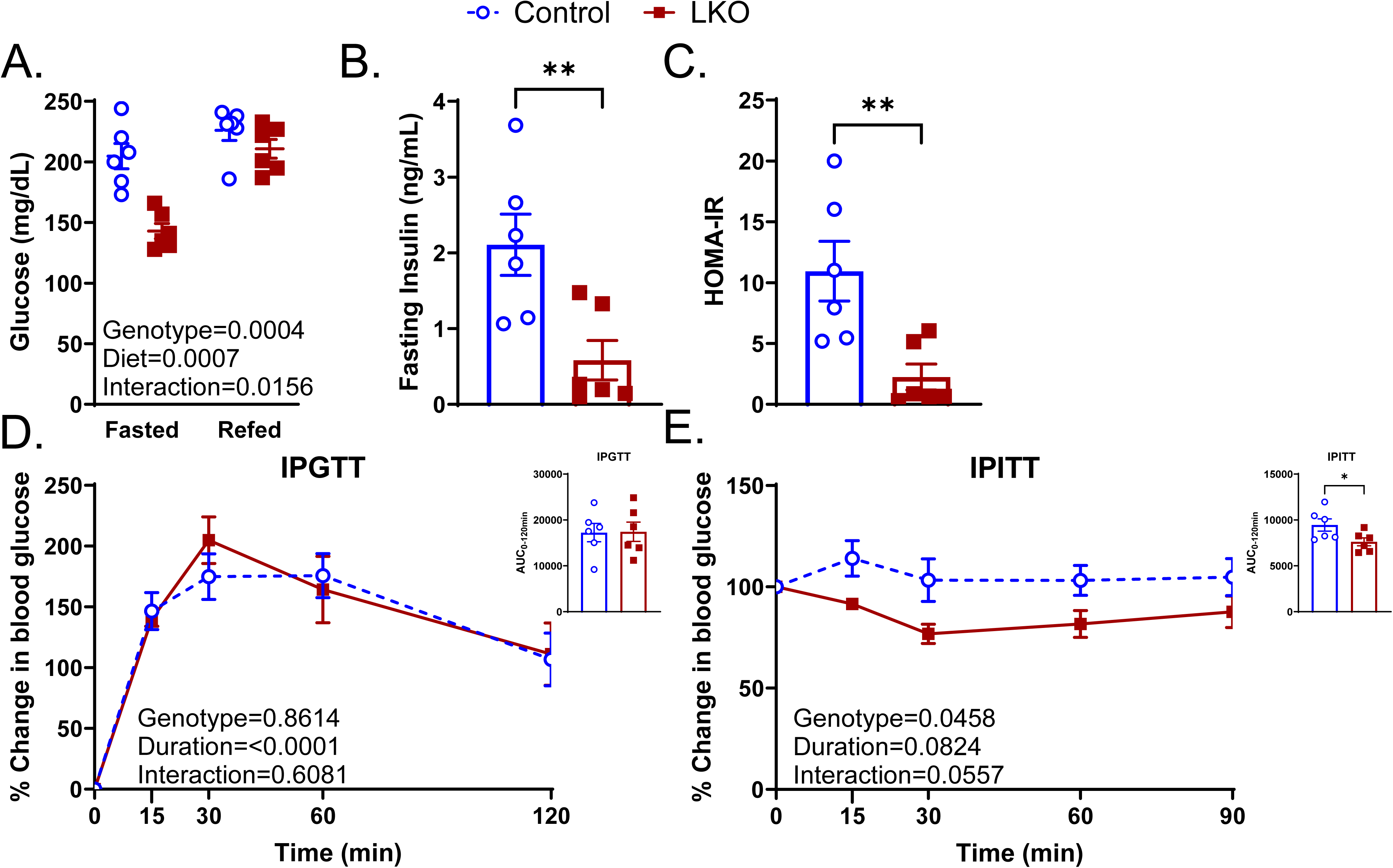
Liver-specific Deletion of Cpt1a Reduces Fasting Insulin and HOMA-IR in Response to HFD-feeding. (**A**) Fed and fasted glucose levels (*n*=6). (**B**) Fasting insulin and HOMA-IR calculations in WT verse LKO mice (*n*=6). (**D, E**) Intraperitoneal glucose (**D**) and insulin (**E**) tolerance tests in WT and LKO mice (*n*=6). The data are presented as % change in blood glucose levels from baseline. The AUCs were calculated and provided in the inset. Significance was determined by a two-way ANOVA with Tukey’s multiple comparison *post hoc* analysis (panel A), a repeated measures ANOVA (D, E), or by a Student’s two-tailed *t*-test as shown in panels B, C, and in the insets. The ANOVA results are provided.

**Supplemental Figure 3.**
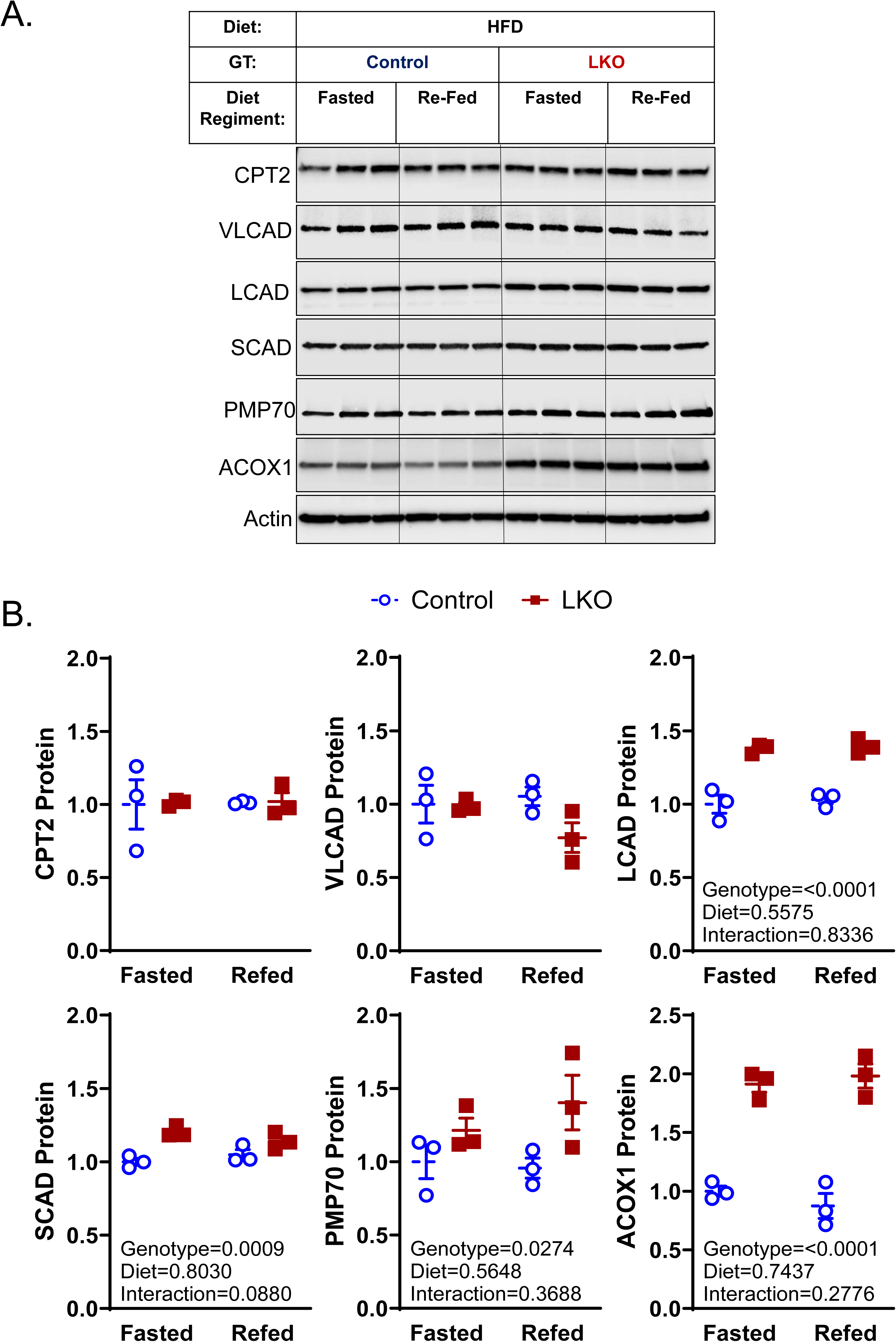
Ablation of Cpt1a in the Liver Increases Mitochondrial and Peroxisomal FAO Protein Levels. (**A**, **B**) Mitochondrial (CPT2, VLCAD, LCAD, SCAD) and peroxisomal (PMP70, ACOX1) fatty acid oxidation proteins were measured by immunoblotting (**A**) and were further analyzed by densitometry (**B**). Significance was determined by two-way ANOVA with Tukey’s multiple comparison *post hoc* analysis (n=3). The ANOVA results are provided.

**Supplemental Figure 4.**
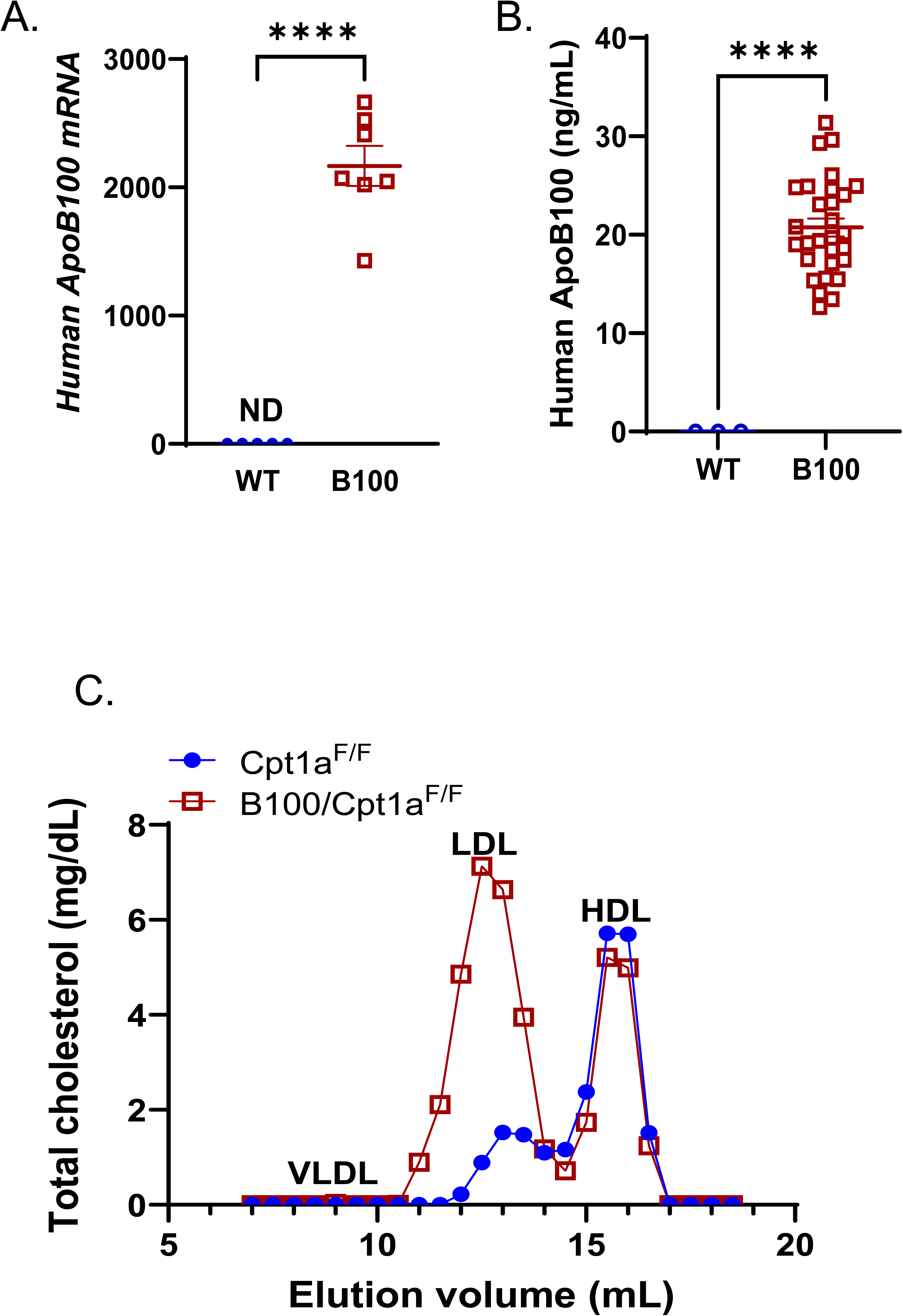
Human B100-transgenic Mice Exhibit a Lipoprotein Profile More Similar to Humans. (**A**) Human ApoB100 RNA levels were measured from livers of WT and B100-Tg mice by QPCR (*n*=5-7). (**B**) Human ApoB100 protein levels were measured in plasma by an ELISA (*n*=3 from WT; 29 from B100-Tg). (**C**) Cholesterol levels were measured from FPLC-fractionated plasma collected from *Cpt1a^F/F^* and B100·*Cpt1a^F/F^* mice. Significance was determined by a Student’s two-tailed *t*-test. ****P<0.0001.

**Supplemental Figure 5.**
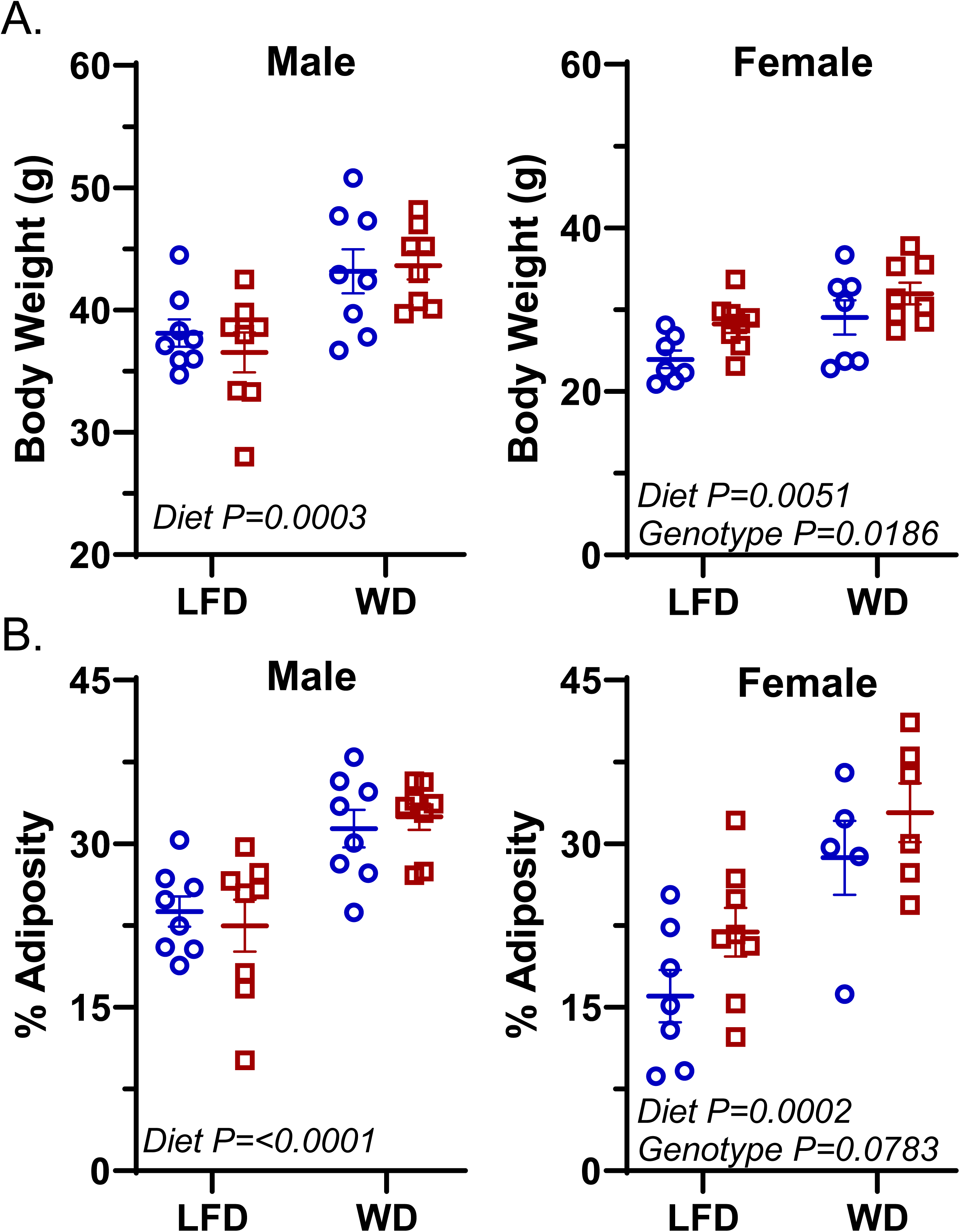
Body Weights nor Adiposity Levels are Altered in B100^Con^ verse B100^LKO^ Mice. (**A**, **B**) Final body weights (**A**) and % adiposity (**B**) levels (e.g. fat mass/total weight) were measured across male and female mice at the conclusion of the experiment. Significance was determined by two-way ANOVA with Tukey’s multiple comparison *post hoc* analysis (*n*=6-8). The ANOVA results are provided.

**Supplemental Figure 6.**
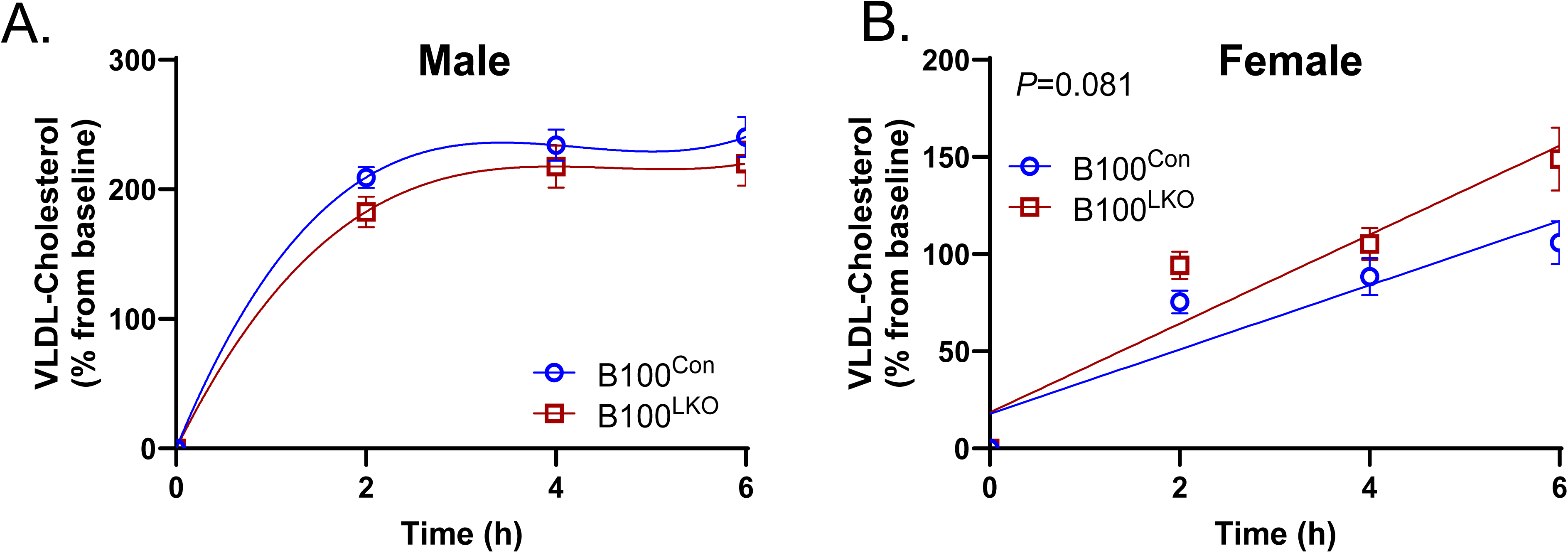
VLDL-C Secretion Tends to Increase in Female B100^LKO^ Mice. (**A**, **B**) Plasma cholesterol levels were measured in male (**A**) and female (**B**) B100^Con^ and B100^LKO^ mice after P407 treatment (*n*=6-11). Nonlinear regression was performed in males while linear regression analyses were performed in females. P-values are provided.

**Supplemental Figure 7.**
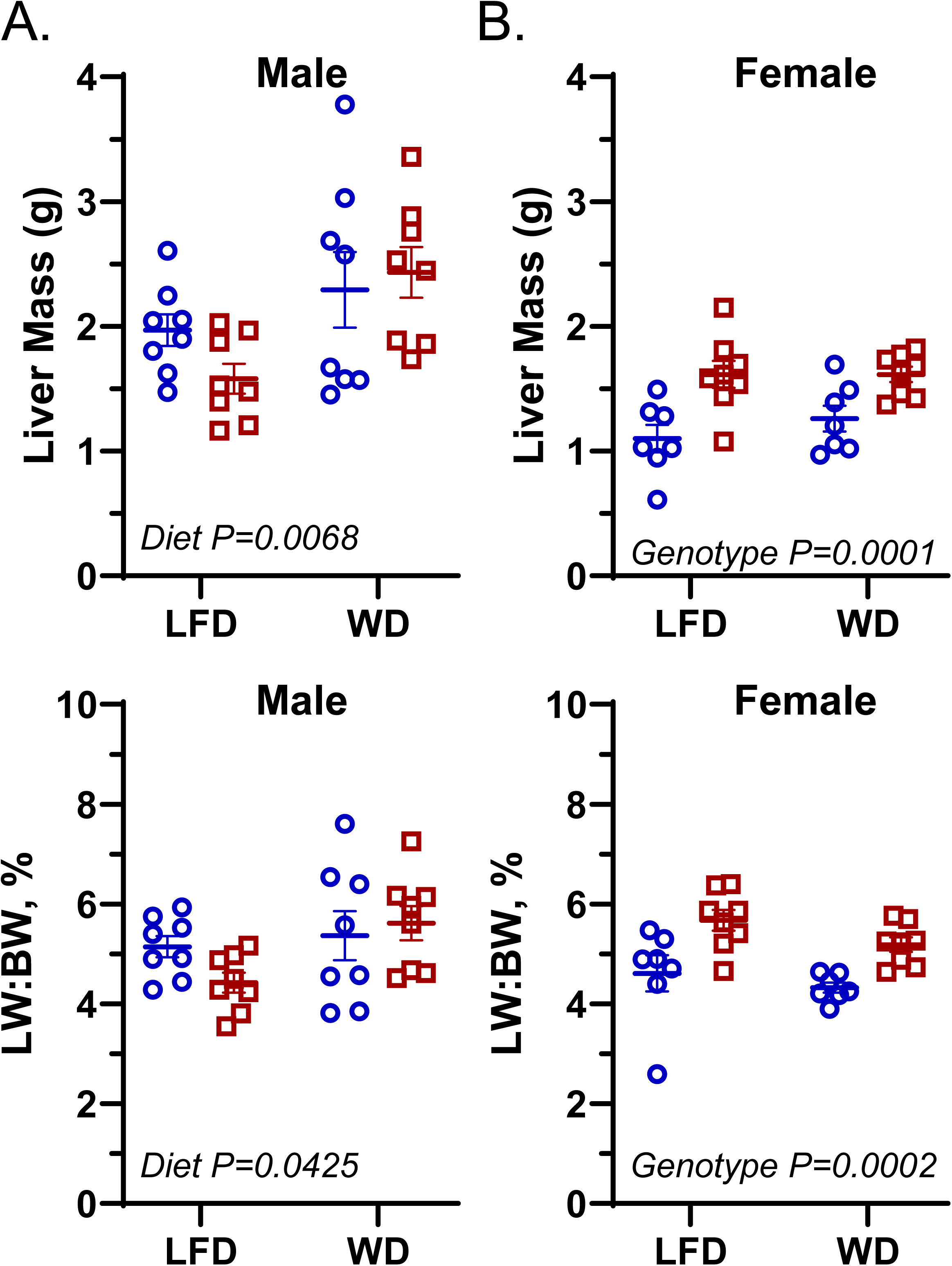
Liver Weights Are Greater in Female B100^LKO^ Mice Independent of Diet. (**A**, **B**) Liver weight and liver weight/body weight ratios in male (**A**) and female (**B**) B100^Con^ and B100^LKO^ mice. Significance was determined by two-way ANOVA with Tukey’s multiple comparison *post hoc* analysis (*n*=6-8). The ANOVA results are provided.

**Supplemental Figure 8.**
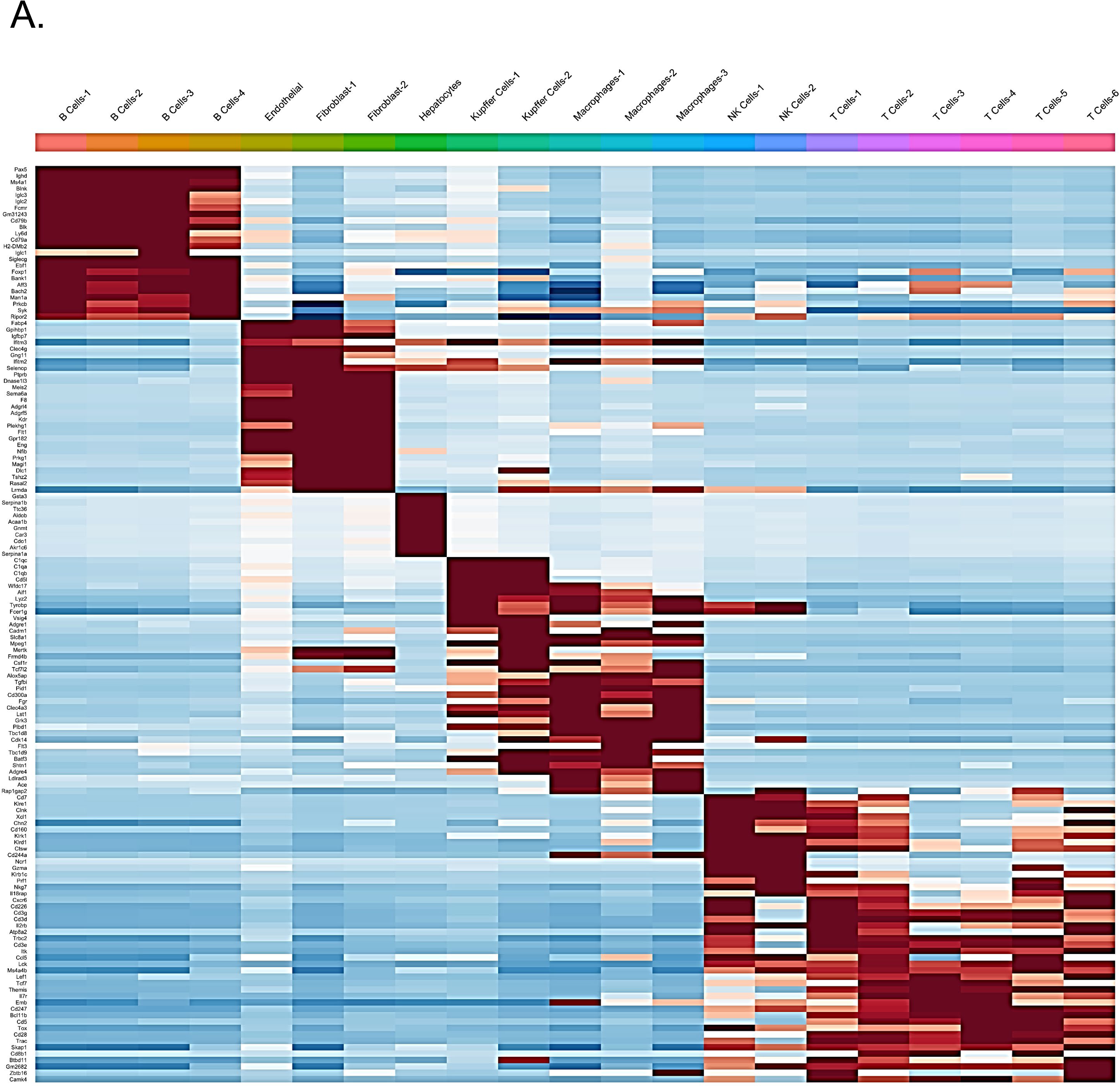
Top Genes Enriched Across 21 Liver Cell Populations. The top genes most highly expressed in each of the 21 cell types from liver scRNA-sequencing data in Figure 8.

